# NeuroGraphBench: Interacting with *Drosophila* Connectomes at Scale for Exploring the Functional Logic of Neural Circuits

**DOI:** 10.64898/2026.08.22.746456

**Authors:** Aurel A. Lazar, Shashwat Shukla, Yiyin Zhou

## Abstract

*Drosophila* connectomic datasets provide increasingly comprehensive maps of neuronal morphology and synaptic connectivity, offering an unprecedented opportunity to explore the structural organization of its neural circuits. This calls for designing automated tools to interact with connectomic datasets at scale for efficiently exploring structural features embedded in the vast amounts of data. Yet the central challenge remains the understanding of the functional logic of neural circuits. In order to formulate a connectome-grounded hypothesis for what a circuit computes, it is critical to (i) characterize the objects in the natural environment in which brain circuits operate, and (ii) formulate how brain circuits represent and process the defined objects. To develop and demonstrate a methodology for these requirements, we focus on the *Drosophila* looming-evoked escape pathway.

We modeled the trajectory of looming objects that are on a collision course (direct-hits) or pass by the fly (near-misses): their projected images on the retina can be characterized by the subtended solid angle and center elevation. By analyzing the sub-neuronal structure and retinotopic organization of the looming pathway embedded in connectomic datasets, we devised a pathway abstraction, and mapped its structural elements into a connectome-grounded functional model. We demonstrate that (i) the expanding boundary of a looming object activates an annulus of LPLC2 neurons, and (ii) the Giant Fiber reads out this population code through its two dendritic branches, modeled as separate units whose outputs are combined to drive its axon. It is this functional separation of the dendritic branches that gives the model access to the center elevation, separating direct-hits from near-misses well before the solid angle of the two trajectories diverge.

To accelerate the connectomic analysis step, we developed a Python toolset with an agentic, code-free workspace interface called NeuroGraphBench (NGB). NGB provides four composable morphology-analysis primitives and an AI Agent that composes them to interactively respond to natural-language queries aided by visualization on an interactive 3D canvas. Thus, NGB automates tedious and repetitive tasks to enable faster and scalable connectomic exploration, keeping human reasoning, instead of writing code, at the center of an open-ended research inquiry.

## 1 Introduction

Connectomic reconstructions now provide increasingly comprehensive maps of neuronal morphology and synaptic connectivity, from single neuropils to whole brains [1, 2, 3, 4], offering an unprecedented opportunity to explore the structural organization of neural circuits. A central challenge is to turn this structure into an understanding of function [5, 6, 7]: how can the structural features embedded in a connectome be efficiently explored, and how can they be used to uncover the functional logic of a brain circuit? Structure alone, however, is not sufficient. The morphological and connectivity features of a circuit do not immediately translate into its functional logic [8, 9]. Characterizing the sensing environment in which the circuit operates is essential in determining its functional logic. To develop and demonstrate our methodology for uncovering the functional logic of a concrete circuit we focus on the *Drosophila* looming-evoked escape pathway.

As illustrated in Figure 1, we first modeled the looming object trajectories in the visual space. To escape an impending collision, a fruit fly must judge from its expanding retinal image whether an approaching object is truly on a collision course, and launch in time if it is. Its escape is driven by the GF, a large descending neuron forming the most direct route from the visual system to the motor centers [10, 11]. The escape getaway comes at a cost, as the wide GF axon that carries it out is metabolically expensive and, in addition, sacrifices the postural stability of the fly: the GF should thus only be triggered by looming objects that will likely strike it [12, 13]. A genuine collision is not evident from the expansion of the retinal image alone, as illustrated in Figure 1a: an object on a direct-hit trajectory (teal) keeps its expansion centered on the fly’s line of sight, whereas one that will narrowly miss (orange) expands as well while its center drifts in elevation. Even though not yet shown in the fruit fly, the locust does discriminate collisions from near-misses with their Descending Contralateral Movement Detector [14]. Yet existing models of and experiments on looming-detection circuits have not fully characterized the geometry of near-miss trajectories: they either ignore near-misses entirely or treat them only as a type of looming input [15, 16, 17]. While *Drosophila*’s response to near-misses has not yet been investigated, it is natural to ask if some functional circuit model can deal with this setting as well.

**Figure 1:**
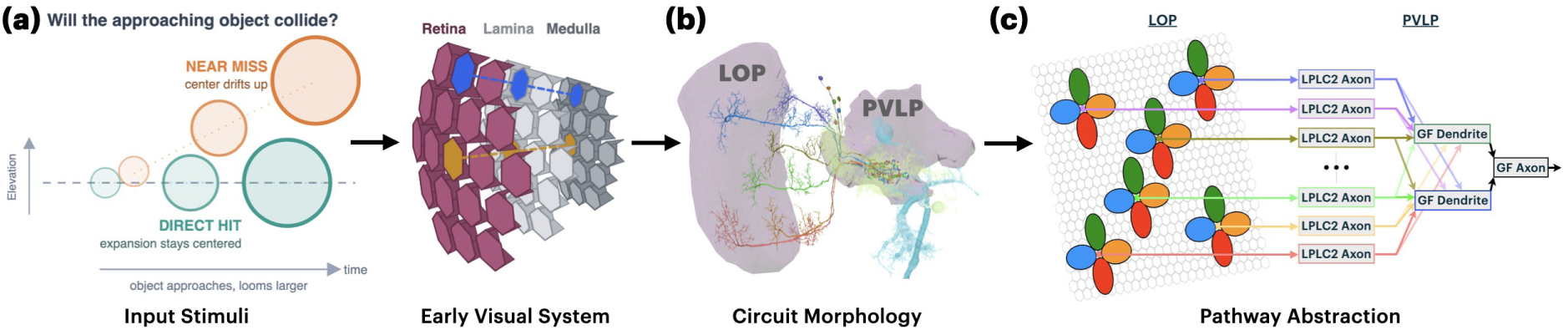
Connectome-grounded modeling of looming-detection and pathway abstractions of the fruit fly brain. **(a)** Left: The *input stimuli* : an object approaching on a direct-hit trajectory (teal) keeps its expansion centered on the elevation of the fly, whereas an object on a near-miss trajectory (orange) drifts in elevation as it approaches the fly’s retina. Right: schematic of the *early visual system* of the fruit fly brain, whose retinotopic organization is carried column by column from the retina, to the lamina and the medulla, illustrated by two example columns (blue and gold). **(b)** The *circuit morphology* of the escape pathway, in which the dendrites of LPLC2 neurons innervate the Lobula Plate (LOP) and their axons converge onto the Giant Fiber (GF) in the Posterior Ventrolateral Protocerebrum (PVLP). **(c)** The *pathway abstraction* constructed from this morphology: each LPLC2 neuron integrates cardinal motion over a four-arm retinotopic receptive field, whose four lobes reflect the colors of the corresponding lobula-plate layer, and projects onto the dendrites of the GF.

In the second stage, depicted in Figure 1b, we model the morphology of the looming circuit and identify its structural features. We analyze the circuit’s morphology in the OpticLobe [3], Hemibrain [1] and MaleCNS [18] datasets, and explore further synaptic connectivity of this pathway in the FlyWire [2] dataset. Furthermore, we go beyond simply retrieving aggregate synaptic connectivity between neurons in two key ways. The first is the retinotopic structure of the motion inputs, that is, how the local motion signals carried by the T4 and T5 cells [19, 20, 21] are arranged across the visual field (inherited from the retinotopic organization of the retina, lamina, and medulla [22, 23, 24]). The second is the connectivity of the neurons’ dendrites and axons, that is, how each LPLC2 neuron’s dendrites are distributed across the layers of the Lobula Plate (LOP) [19] and how its axon connects onto the individual dendritic branches of the GF [25, 16].

After modeling the morphology of the looming detection circuit, we abstract the detailed morphology into the pathway shown in Figure 1c. The pathway abstraction incorporates structural features found in both the LPLC2 and the GF. Here, as indicated by the colors on the left of Figure 1c, we schematically model the LPLC2’s dendritic branches in each of the 4 LOP layers as receptive fields. The population of LPLC2s then provides input onto two dendritic branches of the GF, which we will detail in Section 2. The structural organization of the LPLC2 GF pathway depicted in Figure 1c alone does not explain how the fly distinguishes the two approach trajectories; instead, by assigning a model of computation to each structural element, we turn the abstraction into an executable circuit model. We then test this connectome-grounded candidate model over a set of visual inputs that correspond to direct hit and near-miss trajectories, as seen by the compound fly eye.

Under this workflow, analyzing the morphology of the looming detection circuit is one of the key steps in understanding its connectome-grounded functional logic. To accelerate this step, we developed a Python toolset with an agentic workspace interface called NeuroGraphBench (NGB). NGB provides four composable analysis primitives, and the NGB workspace exposes these primitives through a code-free, natural-language interface in which an AI Agent composes them into analyses and renders the results on a live 3D canvas. This workspace allows Human Explorers to interact with the connectome datasets through both text-based conversation and visual inspection. This makes the flexible modeling of morphology, and hence the discovery of new circuit abstractions, practical.

As the same primitives and code-free interface apply unchanged across datasets, across neuropils, and from the whole brain down to the sub-neuronal level, the proposed workflow also scales: the same analysis can be carried out over the Hemibrain, FlyWire, MaleCNS and OpticLobe datasets, extended from a single neuropil to a brain-wide search over pathways, and refined to the resolution of individual dendritic and axonal branches within a neuropil. The AI Agent handles multi-step tool composition in response to Human Explorer queries, thus automating tedious and repetitive tasks to enable faster and scalable connectomic exploration.

This manuscript is organized as follows. In Section 2 we abstract the structure of the looming escape pathway in the fruit fly brain connectome. We introduce the morphology analysis requirements of the escape pathway and the four NGB primitives that address them, and describe the agentic workspace through which those primitives are composed, and the software architecture that supports them. We then provide a pathway abstraction of the LPLC2 GF escape circuit, and provide demos for the NGB agentic workspace. In Section 3, we describe the trajectory of looming objects, and the functional logic of our connectome-grounded looming detection circuit model. We demonstrate and provide insights on the effectiveness of our model in deciding between direct-hits and near-misses.

In Section 4, we provide a brief discussion of our overall methodology for exploring the connectome-grounded functional logic of the looming detection circuit and more generally, fruit fly brain circuits. We also discuss NGB’s role in the future of connectomic analysis of the *Drosophila* brain, as well as the brains of other organisms.

## 2 Exploring the Connectome of the Looming Detection Path-way with NeuroGraphBench

Section 2.1 establishes the morphology analysis requirements of the escape pathway and the four NeuroGraphBench primitives that meet them. The anatomical features of the looming-detection pathway are summarized in Section 2.2. Section 2.3 describes the agentic workspace through which those primitives are composed, the software architecture that supports them, and how natural language and visual interactions in the workspace enable a Human Explorer interactive and scalable exploration of fruit fly connectomic datasets. Section 2.4 illustrates the exploration of *Drosophila* connectome datasets in the NeuroGraphBench workspace through narrated video demonstrations.

### 2.1 Morphology Analysis Requirements for a Looming-Detection Pathway

As depicted in Figure 2, we present here a set of four morphology analysis primitives, namely, columnar mapping, volumetric segmentation, skeletal segmentation and topographic mapping. The compositions of subsets of these four analysis tools serve as a foundation for a wide range of analyses necessary to reveal insights on brain circuit connectivity from connectomic datasets, at the population and subneuronal resolution level. To demonstrate their value, we use the LPLC2→GF escape pathway shown as an example in Figure 2.

*Columnar mapping* serves two purposes in preserving the retinotopy at each of the visual processing stages. First, it propagates the discrete hexagonal coordinates (see also Figure 1a, right and Figure 2a) assigned to each ommatidium in the Retina, or the corresponding cartridge in the Lamina, or column in the Medulla/Lobula/Lobula Plate to *columnar neurons*. Second, for any neuron (including non-columnar neurons such as LPLC2s) having synaptic connections from/to columnar neurons, it characterizes the neuron’s columnar coverage through its columnar partner’s identity. For each LPLC2 neuron, for example, we obtain columnar input maps for each of the four subtypes (a,b,c,d) of T4s and T5s that each provide direction-selective motion inputs along one of four cardinal directions (front-to-back, back-to-front, up and down, respectively) [26, 19, 27]. We model the right eye throughout, and name directions as they appear in the visual field depicted in Figure 2a. Front-to-back and back-to-front motion therefore appear as leftward and rightward motion (due to the first optic chiasm that invert the anterior-posterior axis), respectively, and for simplicity, we shall refer to the four directions as left, right, up and down in what follows.

*Volumetric segmentation* partitions the arbor of a single neuron by the anatomical regions it innervates, and retrieves the synapses that fall within each region. For the escape pathway of Figure 1b, it lets the LPLC2 pathway to be examined within the LOP and its constituent layers and, separately, within the PVLP (Figure 2d). As shown in Figure 2e, we used volumetric segmentation to verify that the synaptic inputs from each subtype arborize on a distinct branch of the dendrite, with one primary branch innervating each layer of the LOP. This allows us to infer that each dendritic branch separately integrates motion fields along one cardinal direction and these dendritic outputs are subsequently combined. The same volumetric segmentation primitive isolates the LPLC2 axon terminal within the PVLP glomerulus, and we find that each LPLC2 neuron provides synaptic inputs to the GF near the tips of its axonal branches (Figure 2f).

**Figure 2:**
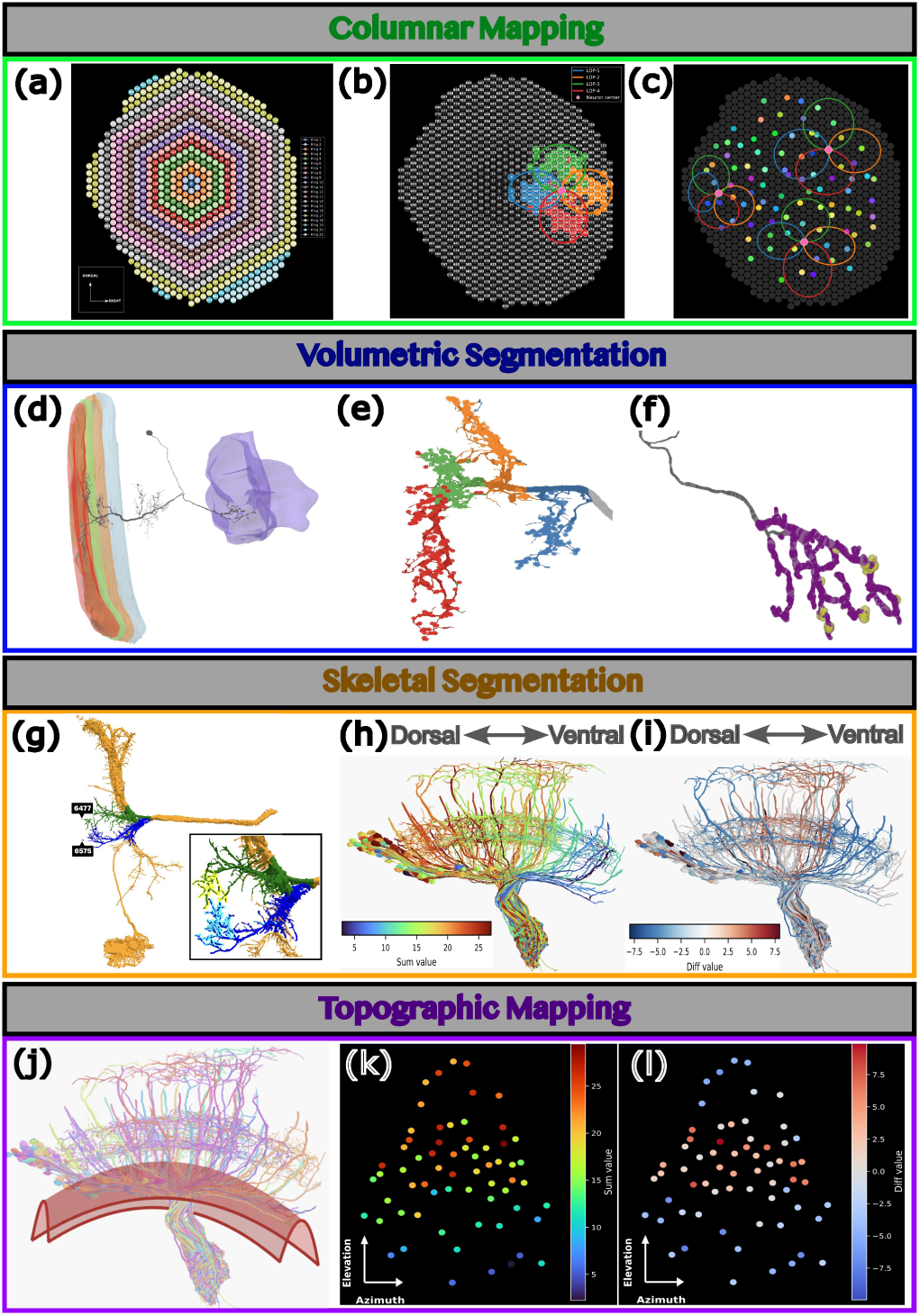
The four morphology analysis primitives. **(a-c) Columnar mapping**. (a) The hexagonal retinal grid that defines the retinotopic coordinate system on which downstream columnar neurons are organized, with concentric rings indicating eccentricity from the center of the grid. (b) Columnar mapping places the columnar T4s and T5s feeding an LPLC2 neuron on the retinal grid, colored by cell subtype (a, b, c, d innervating, respectively, the LOP layers LOP-1 to LOP-4), with the retinotopic center of the LPLC2 marked in pink. (c) The same construction across the population of LPLC2s, showing LPLC2 centroids on the retinal grid with the elliptical dendritic receptive fields overlaid for 3 LPLC2s. **(d-f) Volumetric segmentation**. (d) A single LPLC2 neuron spanning the four lobula-plate layers (colored slabs) and terminating in a glomerulus in PVLP. (e) Volumetric segmentation of the LPLC2 dendrite into its four subunits, each in a different lobula-plate layer. (f) The LPLC2 axon terminal segmented within the PVLP glomerulus (purple), where it provides synaptic inputs (yellow) to the GF. **(g-i) Skeletal segmentation** (g) The GF (orange) with its lateral dendrite partitioned into dorsal (green) and ventral (blue) branches by skeletal segmentation, each retrieved as the local segment within a prescribed geodesic radius of a point selected at the branch tip (black markers); *inset:* the two branches and the location of LPLC2 input synapses on each branch in yellow and cyan, respectively. (h-i) The LPLC2 population morphology, with each neuron colored by (h) the *sum* and (i) the difference of the synapse counts its axon makes onto the two GF branches. **(j-l) Topographic mapping**. (j) LPLC2 axons intersected with a projection manifold placed by the Human Explorer (red), yielding a two-dimensional topographic coordinate for each axon. (k) The axon intersection location with the projection manifold in (j) applied on (h) where neurons are colored by the *sum C*_1_ +*C*_2_ of the synapse counts of that LPLC2, then flattened onto a 2D plane: a monotonic gradient along the dorsoventral axis of the axon bundle. (l) The location of intersection between axon and the same projection manifold applied on (i) where neurons are colored by the *difference C*_1_ *C*_2_ of synapse counts, then flattened onto a 2D plane: a biphasic profile along the dorsoventral axis, with no coherent gradient along the anteroposterior axis.

*Skeletal segmentation* retrieves a local segment of a single arbor from a point the Human Explorer selects on the rendered skeleton: given that point and a prescribed geodesic radius, the primitive returns the connected neurite lying within that geodesic distance along the skeleton, together with the synapses it carries. The dendrite of the GF bifurcates into a dorsal and a ventral branch (see Figure 2g) [16, 28]. To isolate these branches, we select a point at the tip of each branch and prescribe a geodesic radius for each, chosen so that the two retrieved segments span the dorsal and ventral branches and terminate at the base of the axon (black markers, Figure 2g). Assigning each LPLC2 synapse to the branch segment on which it arborizes then resolves how each LPLC2 axon contacts the two branches.

*Topographic mapping* assigns continuous 2D coordinates to a population of neurons, recovering how they are arranged relative to one another [29], when columnar mapping cannot be applied. Columnar mapping, by contrast, anchors those coordinates to the hexagonal grid by following the columnar pathways, and thus directly yields columnar positions rather than relative ones.

The need for topographic mapping is motivated by the fact that while the columnar maps of the LPLC2 population were estimated in the OpticLobe dataset, the GF and synaptic inputs to it from the LPLC2 population are not available in the same dataset. Consequently, we analyzed instead the morphology of LPLC2 axons and their inputs to the GF in the Hemibrain dataset [1]. The latter dataset does not contain the columnar inputs to the LPLC2 dendrites, and therefore, columnar mapping cannot be applied here. To approximate the retinotopic centers of the LPLC2 population in Figure 2j, we used topographic mapping over the parallelly oriented bundle of LPLC2 axons (before they converge onto a densely interconnected glomerulus). The curvature of the projection manifold, as shown in red in Figure 2j, can be interactively configured to be placed where LPLC2 axons still run parallel to each other and to match the observed curvature of the LPLC2 axon bundle, and thereby its intersection with each axon yields a faithful 2D coordinate.

Topographic mapping presents a less precise (due to the need for the Human Explorer visually configuring the projection manifold) but more flexible primitive than columnar mapping, that can be applied without the need for anchoring on columnar neurons. It is required precisely because connectome datasets are frequently incomplete: the Hemibrain dataset reconstructs the Giant Fiber and the LPLC2 axons that contact it, but not the columnar inputs that would fix the retinotopic position of those LPLC2s.

### 2.2 Pathway Abstraction of the Looming Detection Circuit

We used columnar mapping and volumetric segmentation to characterize the connectomic structure of LPLC2 dendrites and their retinotopic motion inputs, and used skeletal segmentation and topographic mapping to explore two readouts from LPLC2 axons by the GF dendrites. Collating these connectomic findings yields the pathway abstraction of Figure 1c. This pathway abstraction serves as the structural substrate for the architecture of the circuit model described in Section 3. Here, we summarize the key features of the pathway abstraction of the LPLC2 GF looming detection circuit.

To characterize the T4/T5 inputs onto each distinct dendritic branch of LPLC2 neurons in the LOP (see Figure 2e), we first applied columnar mapping on T4 and T5 neurons in the OpticLobe dataset [3], the main feedforward inputs onto LPLC2s in the LOP. We assigned a column label to each T4 and T5 neuron, and retrieved the number of T4 and T5 neurons that synapse onto each LPLC2 dendritic branch, together with the respective synaptic counts. Consistent with previous connectomic studies [3], each subtype of T4 and T5, carrying respectively ON and OFF motion signals in one of the four cardinal directions [19] in the same visual area, converge onto the same LPLC2 dendritic branch, with T5 neurons providing slightly more synapses than T4 neurons in general.

The column labels carry over from the T4 and T5 neurons to their postsynaptic targets: the columns of the T4 and T5 neurons synapsing onto a given LPLC2 dendritic branch define the columnar coverage of that branch. The columnar coverage of each of an LPLC2’s dendritic branches can be modeled by an elliptical motion input map, as shown by blue, orange, green and red ellipses in Figure 2b. They emanate radially from a shared retinotopic center at their intersection which we refer to as the center of an LPLC2 neuron (see pink dot in Figure 2b). The centers of all 91 LPLC2 neurons in the OpticLobe dataset are visualized together in Figure 2c. We find that (i) the LPLC2 population tiles the visual space, and (ii) the size and orientation of the elliptical LPLC2 motion input maps is vary only modestly across the population, with noteworthy distortions near the periphery.

In addition, the feedforward pathway described above is also influenced by inhibitory feedback mediated by Lobula Plate intrinsic (LPi) neurons. Four types of LPi neurons, namely LPi12, LPi21, LPi34 and LPi43, are particularly relevant. As the name suggests, each of these LPi types receives aggregated T4/T5 inputs in one LOP layer and feedback onto the T4/T5 neurons that terminate in the LOP layer corresponding to the opposite motion direction. For example, LPi12 neurons receive T4a/T5a inputs from LOP-1 and inhibit T4b and T5b neurons in LOP-2. Additionally, they also synapse onto the LPLC2 dendritic branch in LOP-2. These LPi neurons have been shown to underlie motion opponency [20].

As shown in Figure 2g, the axon of each LPLC2 neuron synapses onto both dendritic branches of the GF. Based on the Hemibrain dataset [1], we identified the individual LPLC2 GF synapses on each of the two GF branches using skeletal segmentation. Denoting the number of synapses from LPLC2 *i* onto the dorsal (Figure 2g, green) and ventral (Figure 2g, blue) GF dendritic branches as, respectively, 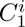 and 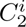, we found no visible pattern of 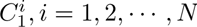, or of 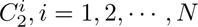 where *N* denotes the total number of LPLC2 neurons. However, we found that the 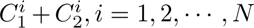 , and the *difference* 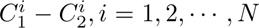 , both had clear spatial patterns. As shown in Figure 2h, coloring each LPLC2 neuron by the sum of its synaptic counts onto the two GF dendritic branches, yields an approximately linear gradient along the ventral-dorsal axis. Likewise, in Figure 2i, the LPLC2 neurons are colored by the difference in synaptic count onto the two branches, yields a biphasic gradient that is positive in the middle and negative at the periphery along the dorsoventral axis. We note that the GF dendritic branches have been described anatomically [16, 28], and while the synaptic gradient corresponding to the sum can be recovered via the typical approach of aggregating synapse counts across both dendrites [25, 30], the novel biphasic synaptic gradient from the difference can only be recovered by estimating synaptic counts onto the two dendritic branches in isolation.

To better visualize and verify the spatial pattern of the sum and difference, we then obtained a topographic mapping of the number of synapses onto each of the GF branches for each LPLC2 neuron. Each LPLC2 axon intersects the projection manifold at a single point, and flattening the manifold onto a plane assigns to that neuron a 2D topographic coordinate, whose axes run along the dorsoventral and the anteroposterior extent of the axon bundle. Plotting every LPLC2 at its topographic coordinate, and coloring it by the sum and difference obtained via skeletal segmentation (Figure 2h, i), yields Figure 2k, l, respectively. We find that the gradients along the dorsoventral axis are preserved in the topographic plane, and that there are no clear gradients along the anteroposterior axis.

To further validate the topographic mapping methodology and the observed spatial profile of the sum and difference of synaptic counts mentioned above, we applied a similar topographic mapping to the LPLC2 axons and a skeletal analysis of GF dendrites in the recently released MaleCNS dataset [18]. We found the same monotonic gradient for the sum and biphasic profile for the difference of synaptic counts (see Figure 9). Because the MaleCNS dataset includes the OpticLobe dataset as a subset, columnar mapping is available for the LPLC2 neurons that synapse onto the GF: we assigned each LPLC2 a center based on it inputs maps from columnar T4s and T5s, without the need for a projection manifold. Figure 9(b,e) plots the sums and the differences of synaptic counts to the GF dendrites at these LPLC2 centers. The linear and biphasic spatial profiles validate those obtained through topographic mapping, both in the MaleCNS (Figure 9(c,f) and Hemibrain (Figure 2k,l) datasets.

In summary, to model the pathway abstraction of the LPLC2 dendrites, each LPLC2 integrates cardinal motion over four elliptical fields extended from its center, and aggregates inputs from both T4 and T5 neurons terminating in the corresponding LOP layer. In addition, LPi neurons receive inputs from T4 and T5 neurons sensitive to motion in one direction and provide feedback onto those sensitive to motion in the opposite direction as well as LPLC2 dendritic branches that integrate the latter. The pathway abstraction at the LPLC2 axon terminal can be characterized by two quantities: the sum and difference of synaptic counts onto the two GF dendritic branches.

### 2.3 Agentic Workspace Design for Connectomic Exploration

Connectomic exploration proceeds as a cycle: exploring a dataset establishes which neurons comprise a pathway, the structure of those neurons is then analyzed, and what the analysis reveals reframes the next question. Carried out by hand, each analysis is a separate script written against the database, and because successive scripts share no state, each one re-derives the neuron sets and re-establishes the geometric anchors that the previous script already found, and the findings accumulate in notes in addition to the code rather than diving deeper in the analysis itself. The resulting cognitive load grows with the number of exploration turns, the size of the circuit under study, and the number of datasets the exploration spans. In practice it is this load, rather than any limitation of the underlying analyses, that constrains the scale at which one can interactively explore *Drosophila* connectomes at scale.

We therefore designed a *code-free* agentic workspace for NeuroGraphBench (Figure 3): the Human Explorer poses questions in natural language and places the geometric anchors that some analyses require (such as a point on a neuron’s skeleton or a projection plane) by direct manipulation of the rendered morphology; an LLM agent answers by invoking the NGB toolset and carries the findings of each turn forward as context for the next.

**Figure 3:**
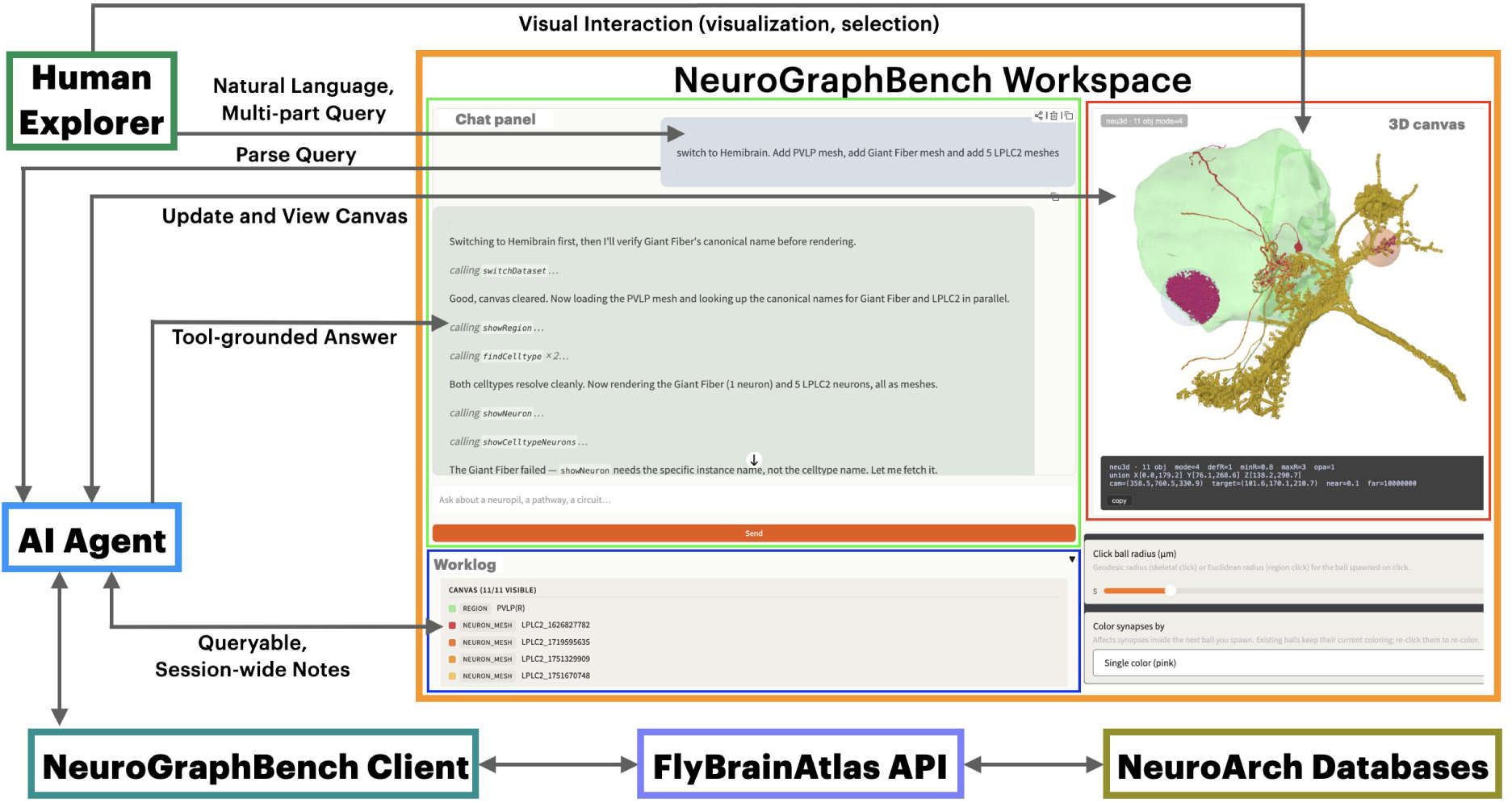
The agentic workspace and the architecture of NeuroGraphBench. The NGB Workspace consists of the chat panel, the worklog, and the 3D canvas. In the chat panel (top left), a multi-part query from the Human Explorer is parsed and answered turn by turn, with the AI Agent naming each NGB function call. The worklog (bottom left) lists the entities currently in play, here the PVLP region and the LPLC2 neuron meshes. The 3D canvas (top right) renders neuropil meshes, neuron morphology and synapses, and provides the controls through which the Human Explorer supplies geometric anchors, such as the radius of the ball spawned on click. The bottom row shows the server-side stack, from the NeuroGraphBench client through the FlyBrainAtlas API to the NeuroArch databases. The *Human Explorer* poses natural-language, multi-part questions, and interacts visually with the 3D canvas through visualization and selection. The *AI Agent* returns tool-grounded answers, updates and views the canvas, and writes queryable, session-wide notes to the worklog.

The workspace comprises an interface (Figure 3, orange box) and a server-side stack (Figure 3, bottom). The Human Explorer and the AI Agent act on the same workspace session through different channels: the Human Explorer through the rendered panels, posing queries and supplying geometric input by direct manipulation; the AI Agent programmatically, parsing each query into typed calls to the server-side stack and accordingly updating the information rendered on the panels.

The server-side stack is built on top of the NeuroArch database [6], which stores the morphology of neuropils and neurons, the locations of synapses, and neurotransmitter information under a unified schema shared across connectome datasets [1, 2, 3, 4]. The server-side stack consists of the FlyBrainAtlas API, a server that exposes the NeuroArch datasets and caches the derived quantities on which the primitives depend, and the NeuroGraphBench Client that presents the four analysis primitives (Section 2.1) and higher-level *composer* functions to an LLM agent. The latter agent translates natural-language questions into a chain of typed tool calls in a single turn, allowing the use of the four analysis primitives without writing a single line of code.

The workspace interface presents the Human Explorer with three panels: a chat panel carrying the dialogue with the AI Agent, a 3D canvas on which the Human Explorer visually supplies input, and a worklog panel that displays the persistent *workspace state*: after each turn the AI Agent records the entities it resolved, the operations it invoked and the findings they produced, so that both the Human Explorer and the AI Agent can query them in later turns. The implementation of these panels and the flow of information between them is described in the Appendix A.1.

Combining natural language conversations with real-time visualizations, in a number of ways, a powerful interface for interacting with connectomic datasets. First, exploration of a dataset is enriched by interactive visualization: the 3D canvas responds to Human Explorer queries visually and in real time, narrated by the natural language dialog. A question such as “Which LOP layers does the LPLC2 dendrite innervate, and where do its inputs sit on the columnar grid?” can be immediately visualized, while the chat panel lists the relevant LOP layers and column numbers, which respond to mouse hover by highlighting the corresponding region of the 3D canvas. Second, supported by natural language dialog, visualization is no longer static. This presents a completely different working experience from the usual code-driven visualization workflow. Human Explorers can immediately ask follow-up questions prompted by new observations of the 3D canvas, without having to work out what code would be needed to perform the analysis. In addition, through *visual selection* on the 3D canvas, the Human Explorer can now supply geometric anchors directly to the dialog. These interactions keep the Human Explorer focused on what matters scientifically, rather than on the mechanics of how analyses must be carried out.

*Scaling* is the payoff: the same code-free workspace lets an analysis grow without the Human Explorer’s effort growing with it, in three ways. (a) The agent absorbs the per-turn bookkeeping that unaided exploration accumulates, namely schema translation, tool discovery, entity tracking, and recall of earlier results, and it invokes *composer* functions, which are codified workflows that chain the primitives into common multi-step analyses, directly rather than recomposing the same primitive chain each time; within a fixed time budget the researcher therefore does more (see also Section 2.4). (b) Because the primitives operate over multiple connectomic datasets [3, 1, 2] through the common NeuroArch database schema, with cached naming conventions and cell-type aliases from the FlyBase ontology [31], the same query automatically acts upon multiple datasets. (c) Composing the primitives lets one analysis reach across the three spatial scales: skeletal segmentation and volumetric segmentation together resolve the *sub-neuronal* scale of a single neurite; volumetric segmentation with topographic and columnar mapping resolves the *neuropil* scale of a population. Since volumetric segmentation and columnar mapping require no visual input, they can be composed across single multi-part natural-language questions, and by chaining from one neuropil to the next they can reach the *brain* scale.

*Pathway search*, a brain-scale workflow that traces geometry-annotated pathways across neuropils and datasets, complements the sub-neuronal depth of the primitives with the breadth of brain-scale exploration.

### 2.4 Exploring *Drosophila* Connectomes with NGB Agentic Workspace

We provide five demonstrations to illustrate how agentic workspace help scale up access to a variety of NGB capabilities.

In Demo 1 (Supplementary Video S1), the Human Explorer volumetrically segments a single LPLC2 neuron in two ways: by the LOP layers it innervates, and subsequently adding synapses on one of the obtained segments; and by visually placing a 3D ball on the canvas, then retrieving the LPLC2’s synapses inside this ball and coloring them by neurotransmitter type. The ball’s center and radius are configured by visual manipulation on the morphology canvas, segment names are accessible by clicking them on the canvas, subsequent segmentation and coloring operations are translated by the AI Agent in the chat panel.

In Demo 2 (Supplementary Video S2), the Human Explorer visualizes the columnar maps of an LPLC2 grouped by its T4 and T5 input subtypes, along with the neuron’s center; the columnar maps of an LPi34 neuron that synapses on this LPLC2; and finally the centers of all LPLC2s at once. No visual operations are required: the AI Agent handles columnar mapping operations in the chat panel, and thus the analysis automatically scales from one neuron to the whole population.

In Demo 3 (Supplementary Video S3), the Human Explorer creates a projection manifold and aligns it with respect to the arbors of a handful of LPLC2s, and projects these LPLC2 arbors onto the manifold. The manifold cannot be specified in words and is configured visually in the morphology canvas, its name copied by visual selection, and the topographic mapping operation is handled by the AI Agent in the chat panel.

In Demo 4 (Supplementary Video S4), the Human Explorer segments out two GF dendritic branches by creating geodesic neighborhoods along the GF skeleton, then colors upstream LPLC2s by the sum, and by the difference, of their synaptic counts onto the two GF branches. The centers and radii for the geodesic neighborhoods are configured visually on the morphology canvas, while the LPLC2 population coloring request is handled by the AI Agent in the chat panel.

In Demo 5 (Supplementary Video S5), the Human Explorer traces feedforward pathways from the photoreceptors to LPLC2 cells in the FlyWire dataset, matches the cell-types the search surfaces in the OpticLobe dataset based on the pathways’ connectivity patterns, and then summarizes the layer-wise innervation and connectivity of every LPi cell type in the OpticLobe. All operations are handled by the AI Agent in the chat panel, wherein the AI Agent also manages the context required to perform these operations (surfaced cell-type names) and summarizes the obtained results (cell-type connectivity patterns across two datasets).

In summary, we demonstrated how the NGB workspace allows connectomic exploration to scale up by providing two code-free interaction methods (natural language prompts and visual morphology manipulation) that alleviate the effort associated with manual exploration. As the AI Agent composes operations and carries the exploration state forward, the per-turn effort stays roughly constant across a conversation, and thus the Human Explorer’s attention stays on the scientific question rather than the mechanics of the exploration.

## 3 Connectome-Grounded Functional Logic of the Looming Detection Circuit

The pathway abstraction of the LPLC2 GF circuit established in Section 2.2 leads in this section into a connectome-grounded circuit model. Section 3.1 assigns a computational role to each structural element of the pathway abstraction, yielding an executable circuit model of collision detection whose putative functional logic is described stage by stage. Section 3.2 then presents the response of our model to direct-hit and near-miss trajectories, evaluates our connectome-grounded functional model and provides insights on how direct-hits and near-misses can be distinguished.

### 3.1 The Functional Logic of the Looming Detection Circuit Model

An approaching spherical object with physical radius *r*, moving at constant velocity *v* and at distance *d*(*t*) from the retina of the fruit fly, subtends on the spherical eye (with unit radius) a retinal disk, denoted as *D*(*t*), with an angular diameter (see also Figure 10)

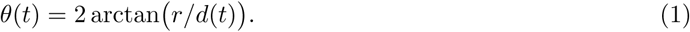

The center of this retinal disk is positioned at elevation *ψ*(*t*). The area of the retinal disk, the solid angle Ω(*t*) = 2*π* 1 cos(*θ*(*t*)*/*2) , is a function of *θ* alone. We consider two trajectories, both beginning on the fly’s line of sight (Figure 1a and Figure 8, Appendix A.2): a *direct-hit*, which stays at centered elevation (*ψ*(*t*) 0) as the distance *d*(*t*) shrinks and the angular diameter *θ*(*t*) grows as the object moves towards collision; and a *near-miss*, which may carry a growth in *θ*(*t*) but drifts off-axis, so that its center of elevation *ψ*(*t*) monotonically rises (or falls) as the object passes by the fruit fly.

Based on the pathway abstraction of the LPLC2 GF circuit, in our connectome-grounded model, the population of LPLC2 neurons tile the visual field. Each LPLC2 radially integrates the local motion field carried by the direction-selective T4 and T5 cells along with opponent motion signals from inhibitory LPi cells (see Figure 4a). As we will show later in this section and in the Appendix C, only the LPLC2s whose receptive fields intersect the expanding boundary of the solid angle are activated, forming an annulus that contains its boundary.

**Figure 4:**
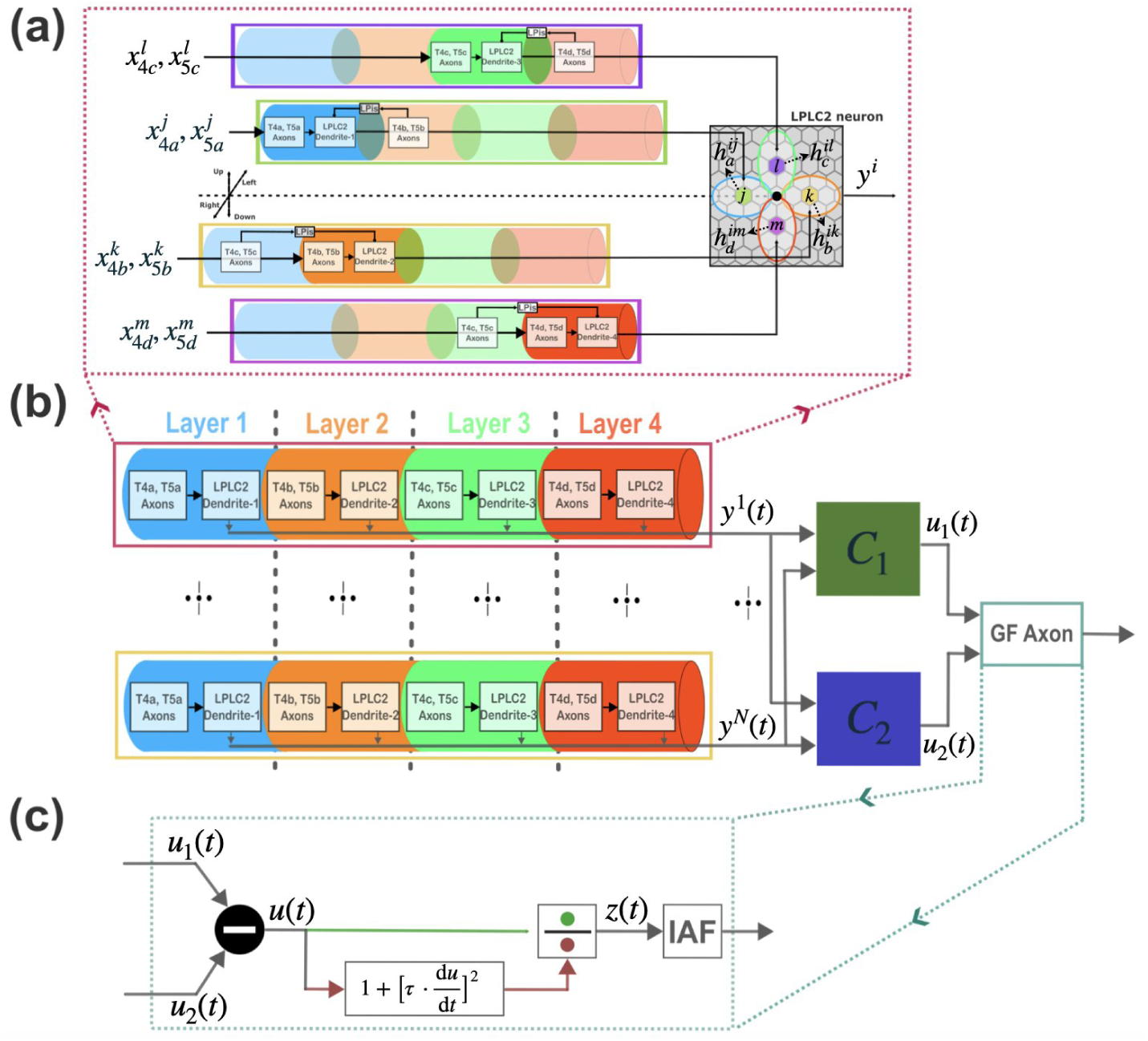
The connectome-grounded looming circuit model. The looming circuit model instantiates the pathway abstraction of Figure 1c, shown at the scale of a single LPLC2 neuron in (a), at the population processing level of the GF dendrites in (b), and at the GF SIZ level in (c). **(a) Model of an LPLC2 neuron.** (left) Each of the four dendritic branches (LPLC2 Dendrite-1 to -4) innervates one LOP layer, shown as the colored bands (blue, orange, green, red), where it receives retinotopic inputs from T4,T5 axons innervating that layer (T4a,T5a to T4d,T5d), each conveying local motion along one cardinal direction (left, right, up, down). In each column, the T4 and T5 axons carrying the opposite cardinal direction drive a LOP intrinsic neuron (LPi), which inhibits that same column’s input to the LPLC2 dendritic branch, so that each branch is driven by the difference between inputs that are sensitive to motion in opposite directions. The LPi’s are colored by the layers in which they receive excitatory inputs and provide inhibitory outputs, shown as from/to LPi connections. (right) The four subunits’ elliptical columnar coverages are modeled as spatial receptive fields. For an LPLC2 neuron *i*, its direction-selective inputs in columns *j, k, l, m* from the T4 and T5 cells are, respectively, 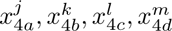 and 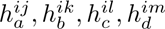. The corresponding receptive field entries are 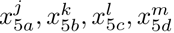, and the axonal output *y^i^*(*t*) is the rectified sum (not shown) of the four dendritic filter responses. **(b) LPLC2 population to GF processing model.** The population of *N* axonal outputs *{y^i^*(*t*)*}^N^* , each layered by LOP depth as in (a), is read by the two dendritic branches of the GF through the synaptic weight vectors **C**_1_ (dorsal) and **C**_2_ (ventral), corresponding respectively to the green and blue branches in Figure 2g, with respective branch outputs *u*_1_(*t*) and *u*_2_(*t*). These dendritic branch outputs are passed to the GF Spike Initiation Zone (SIZ). **(c) Model of the GF SIZ.** The SIZ feeds the difference *u*(*t*) = *u*_1_(*t*) *u*_2_(*t*) of the two dendritic branch outputs to a divisive-normalization processor, whose numerator is *u*(*t*) and whose denominator is set by its rate of change *u̇*(*t*) (the d*/*d*t* and squaring blocks). The decision signal *z*(*t*) drives the ideal integrate-and-fire (IAF) neuron whose spike initiates the escape.

The GF then reads this population code as the difference between its two dendritic branch outputs (see Figure 4b). Its Spike Initiation Zone (SIZ), is modeled as a divisive-normalization processor (DNP) followed by an ideal integrate-and-fire (IAF) neuron that generates the escape spike (see Figure 4c). We describe, stage by stage, the functional logic of the looming detection circuit depicted in Figure 4. For the theoretical grounding details, we refer the reader to the Appendix B.

#### Local Looming Detection by Individual LPLC2 Neurons

In our model, estimates of local motion [21, 32], along the four cardinal directions (left, right, up, down) are carried by the T4 and T5 cell types (a, b, c, d) that each innervate one LOP layer. The axonal outputs of the T4 and T5 cell types in column *j*, *j* = 1*, . . . , M,* are respectively denoted as 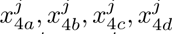 and 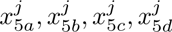, and their pairwise sum per cell type (a, b, c, d) are denoted as 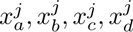 . Each LPLC2 neuron *i*, *i* = 1*, . . . , N,* reads these motion inputs 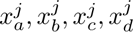 across the columns *j* = 1*, . . . , M* , through four dendritic branches that are modeled as elliptical spatial receptive fields. They are denoted, respectively, as 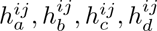, where the superscript indicates the influence of the inputs from column *j* on the LPLC2 *i*. Each of the 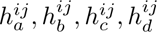 takes the value 1, if the LPLC2 neuron *i* receives inputs from the respective T4/T5 cell types in column *j*, and is 0, otherwise. As schematically depicted in Figure 4a, each receptive field pools the motion inputs from the T4 and T5 cells whose preferred direction points radially outward with respect to the center *P ^i^* of the LPLC2 neuron *i*, and subtracts from them the inputs of the T4 and T5 cells tuned to the opposite, radially inward direction. The subtracted, radially inward inputs reach the branch as inhibition: in each column, an LPi neuron driven by the T4 and T5 cells tuned to the inward direction inhibits that same column’s input to the LPLC2 dendritic branch (see LPi blocks in the tubes in Figure 4a). We note that even though the LPi’s columnar coverage is large, we model the LPi neurons as providing motion opponency column by column. It has been shown that some wide-field neurons, such as the CT1 neuron whose dendrites cover the entire visual field, can be divided into independent functional units in each column [33]. Therefore, we do not model the spatial extent of the LPi’s receptive fields.

The four filtered subunit signals are summed and then rectified to produce the axonal drive *y^i^*(*t*) of LPLC2 neuron, *i* = 1*, . . . , N* , given by

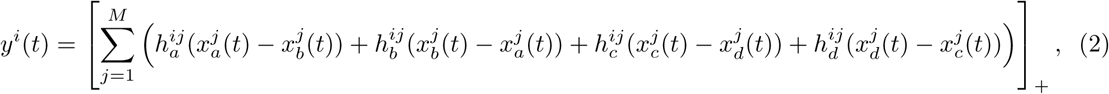

where [ ]_+_ denotes half-wave rectification (for more information, see the Appendix B). Because the summation precedes the rectification, the bracketed sum measures the net outward (expanding) motion over the neuron’s receptive field. Consequently across an approaching object, the LPLC2 neurons whose centers fall in the *interior* of the object’s retinal disk *D*(*t*) receive radially expanding motion inputs in at least one of their dendritic branches (see Figure 12a, top right inset). This will result in a positive value in Eq. (2) before the rectification, and will activate these LPLC2s. For those LPLC2 neurons whose center fall outside the expanding disk, the expanding boundary crosses dendritic branches of these LPLC2s against their preferred directions, thus inhibiting these LPLC2s (see Figure 12a, lower left inset). There is a net negative value in Eq. (2) before the rectification, and these LPLC2s remain silent. Therefore, only those LPLC2s whose centers are inside the retinal disk *D*(*t*) will be activated by the looming object.

#### The Activated LPLC2 Annulus Provides an Estimate of the Solid Angle

In our connectome-grounded model, the *N* LPLC2 neurons uniformly tile the visual field. A looming object of uniform luminance generates local motion only at the boundary of its retinal disk *D*(*t*). Thus, only the LPLC2s whose dendritic receptive fields intersect the expanding boundary of the solid angle receive motion inputs. Of these, the neurons whose centers are located inside the solid angle are activated since the expanding boundary matches the preferred direction of the receptive fields it intersects, resulting in positive responses. For those whose centers are located outside, the expanding boundary moves against the preferred direction of the receptive fields it intersects. This results in negative outputs from these receptive fields, and these LPLC2 neurons remain silent. The active population is therefore an annulus that grows outward around the boundary (see also Figure 12a).

The LPLC2 activation annulus thus represents the expanding boundary of the solid angle: its radius is the angular radius *θ*(*t*)*/*2, and the centroid of the annulus of active neurons is the elevation *ψ*(*t*) from the solid angle’s center. Both the subtended solid angle Ω(*t*) and the center elevation *ψ*(*t*) can be recovered accurately from annulus population activity, as we show in Appendix C (Figure 12b).

#### Dendritic Readouts and Axonal Processing of the Giant Fiber

The GF is modeled as sampling the solid-angle representation through the two branches of its lateral dendrites, whose synaptic weight vectors **C**_1_ (dorsal) and **C**_2_ (ventral) read the population *{y^i^*(*t*)*}^N^* into the two dendritic outputs *u*_1_(*t*) and *u*_2_(*t*) (Figure 4b). Their difference *u*(*t*) = *u*_1_(*t*) *u*_2_(*t*) is the single signal the SIZ acts on. The SIZ model converts *u*(*t*) into an escape decision through a divisive-normalization peak detector (Figure 4c), whose output

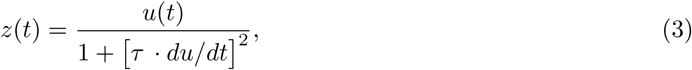

where *τ* is a constant, drives an ideal integrate-and-fire (IAF) neuron (with threshold *δ*) modeling the output of the GF SIZ (see also Appendix B). The IAF neuron emits a spike when the potential reaches the threshold value *δ*, signaling the initiation of the short-mode escape response that leads to a rapid upward launch powered by the mesothoracic legs [13, 10, 34].

#### Functional Logic

The differential readout *u*(*t*) is the result of weighting the active LPLC2 population by the biphasic profile **C_1_ C_2_**, which is positive over a band of elevations centered around the fly’s line of sight and negative above and below it (see Figure 11, Appendix B). While the expanding retinal disk *D*(*t*) lies within the positive band, *u*(*t*) is positive and increases with the solid angle of the object’s retinal projection. Once the disk expands past the band and into the negative flanks, *u*(*t*) begins to decrease and subsequently becomes negative. The readout therefore rises to a positive peak before decreasing.

The direct-hit/near-miss discrimination is developed in detail in the Section 3.2 below.

### 3.2 The LPLC2 Activation Annulus and Its Giant Fiber Readouts

In Figure 5, we computationally evaluate the connectome-grounded model described in Section 3.1 on a direct-hit and a near-miss trajectory, for a looming object that approaches the fruit fly at the same velocity *v* = 0.20 m/s. Our goal is to first explain why our model fires a spike for the direct-hit but not for the near-miss.

**Figure 5:**
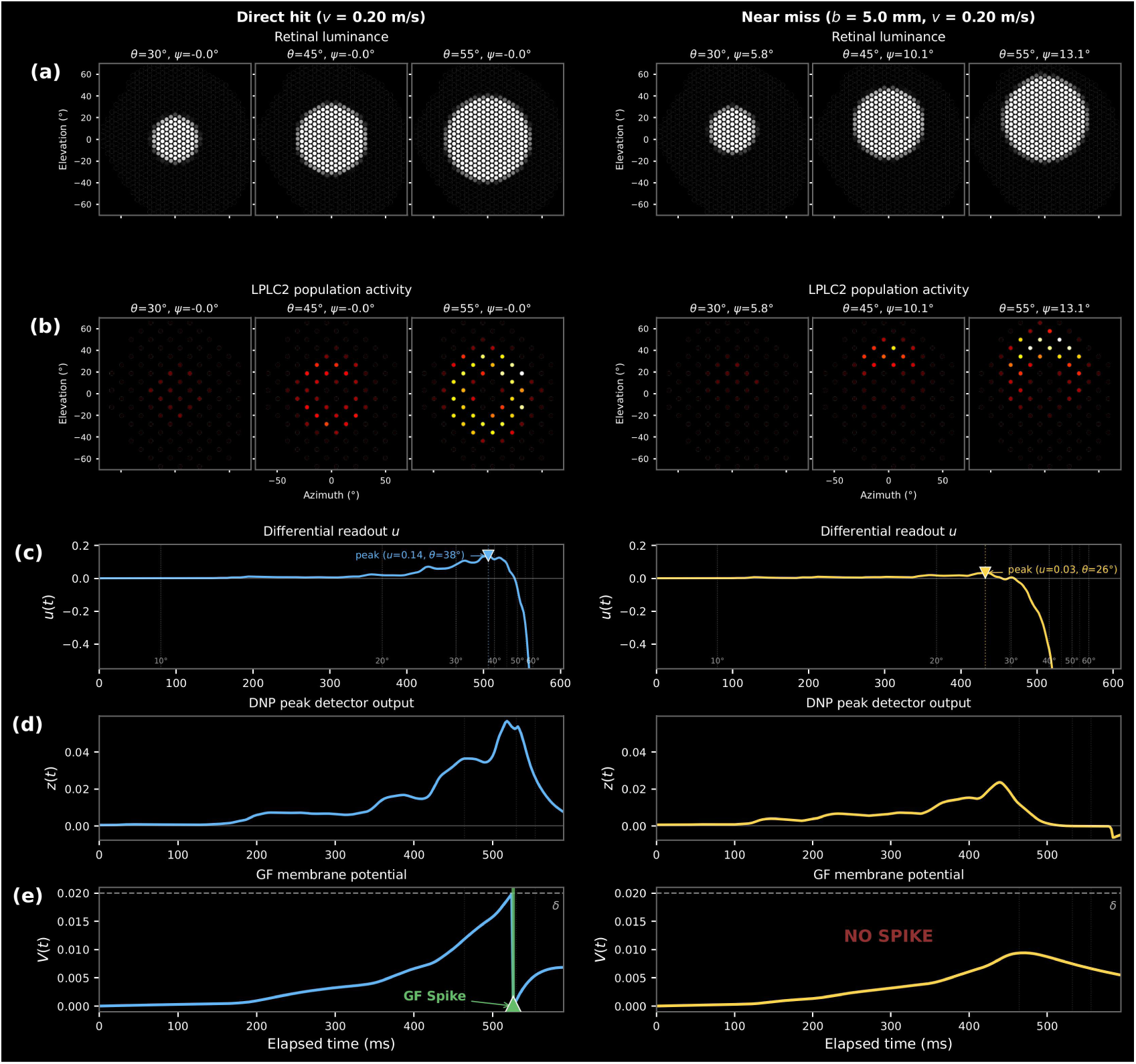
Model response to a direct-hit and near-miss trajectory. Left column: direct-hit (*v* = 0.20 m/s, centered). Right column: near-miss (*b* = 5.0 mm, *v* = 0.20 m/s). **(a)** Retinal luminance at three representative angular diameter values *θ* = 30*^◦^,* 45*^◦^,* 55*^◦^*. **(b)** LPLC2 population activity at the same three angular diameters as in (a); the active neurons tile the retinotopic disk the object subtends, centered for the direct-hit and displaced dorsally for the near-miss. **(c)** Differential readout *u*(*t*): for the direct-hit it rises to a high peak when the object’s angular diameter has grown to *θ* = 38*^◦^*; for the near-miss it peaks lower and at the smaller angular diameter *θ* = 26*^◦^*, then becomes increasingly negative as the active disk leaves the positive band of **C**_1_ **C**_2_. **(d)** Divisive-normalization peak-detector output *z*(*t*): a large and late peak for the direct-hit, a smaller and earlier peak for the near-miss. **(e)** GF membrane potential: it reaches the threshold *δ* = 0.02 only for the direct-hit, producing an escape spike at *θ* = 40*^◦^*, and stays subthreshold for the near-miss.

The number of LPLC2 neurons in the activated annulus grows with the angular diameter of the looming object along both trajectories (Figure 5a,b). This number reflects the solid angle the object subtends and, on its own, cannot separate a direct-hit from a near-miss. The differential readout *u*(*t*), which weights the active LPLC2 population 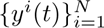 (Figure 5b) by the biphasic profile **C**_1_ *−* **C**_2_ does separate the two (Figure 5c): *u* rises while the activated annulus lies within the positive band of **C**_1_ **C**_2_ and falls once the solid angle expands into the negative flanks. For the direct-hit (Figure 5c (left)), the elevation of the object’s center stays fixed on the line of sight, so the disk grows while remaining centered in the positive band and *u* rises to a high, late peak (*u* = 0.14*, θ* = 38*^◦^*). For the near-miss (Figure 5c (right)), the elevation of the object’s center rises, carrying the growing active disk into the negative flanks earlier, so *u* peaks lower and at a smaller angular diameter (*u* = 0.03*, θ* = 26*^◦^*), then reverses sign as the disk passes fully into the negative band.

The GF SIZ converts this readout into a decision (Figure 5d,e). Its divisive-normalization processor produces an output *z*(*t*) whose amplitude tracks the peak of *u* (compare Figure 5d and Figure 5c), and this amplitude is large for the direct-hit but smaller, and reached earlier, for the near-miss. The ideal IAF neuron that follows accumulates a larger drive for the direct-hit than for the near-miss, and its threshold can be set so that it fires for the direct-hit but not for the near-miss (Figure 5e).

While the stimulus pair chosen above illustrates the mechanism, a fly encounters direct-hits and near-misses across a range of approach velocities, and near-misses across a range of pass-by distances. Its predators span a wide range of attack velocities, from damselflies at around 0.17 m/s [12], through robber flies at 0.5 to 0.9 m/s [35], to dragonflies at 1.4 to 3.75 m/s [36]. The Giant Fiber must launch a timely escape from a direct collision by firing while the time-to-collision still exceeds a takeoff deadline [13]. It must also withhold the metabolically costly escape for an object that will merely pass by [12].

These two conflicting goals, escaping in time and withholding the escape, impose a tradeoff: to escape a fast direct-hit the fly must commit early, for example, by lowering the firing threshold, when a near-miss still looks much like a direct-hit as its center has not yet drifted far and its subtended solid angle is still growing. To illustrate this tradeoff we choose the firing threshold (*δ* = 0.02) for our model such that the GF fires in time to escape all direct-hits with approach velocities less than or equal to *v*_max_ = 0.5 m/s. Note that a maximum escapable looming object approach velocity *v*_max_ necessarily exists. Here the chosen value of *v*_max_ lies in the regime of damselfly approach velocities that fruit flies reliably escape from, and is lower than the faster robber-fly approach velocities [35]. Other choices of *v*_max_ values can be similarly examined.

Figure 6a shows the DNP output *z*(*t*), subtended solid angle Ω(*t*) and center elevation *ψ*(*t*) for a direct-hit (*b* = 0 mm) and near-misses at various pass-by distances (*b* = 2, 5, 10, 20, 40 mm), for a looming object approaching at *v* = 0.3 m/s. For the direct-hit, the solid angle increases over time while the center-elevation remains zero (blue curve). For the near-misses, the solid angle peaks and then falls again while the center-elevation keeps rising. The peak value of *z*(*t*) is much higher for the direct-hit and is progressively smaller for near-misses with larger pass-by distances.

**Figure 6:**
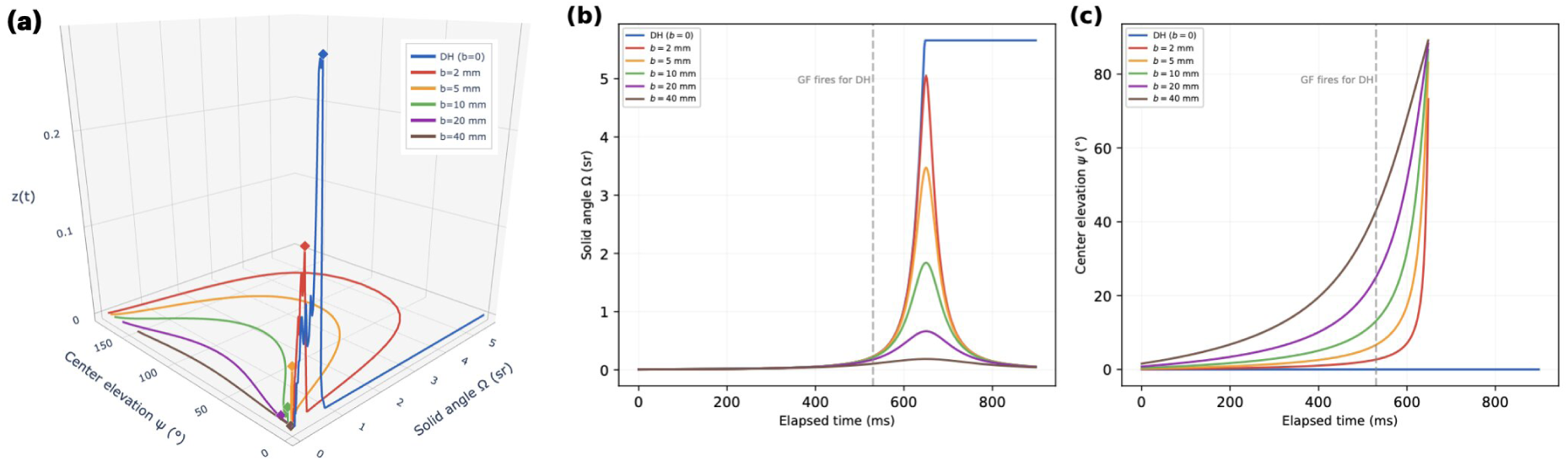
Model dynamics for direct-hit and near-miss trajectories. **(a)** DNP output *z*(*t*) for a direct-hit (*b* = 0 mm) and near-misses (*b* = 2, 5, 10, 20, 40 mm) with approach speed *v* = 0.30 m/s, visualized along the corresponding trajectories defined by the subtended solid angle Ω(*t*) and center elevation *ψ*(*t*). The diamonds indicate the peak response of *z*(*t*), where the peak value decreases with pass-by distance *b*. **(b)** Solid angle Ω(*t*) subtended by the looming object as a function of time for a direct-hit (*b* = 0) and near-misses at several pass-by distances, all at *v* = 0.30 m/s. The solid angle for the direct-hit increases monotonically till the object expands to cover the entire screen, while the solid angle rises, attains a peak and then falls for near-misses. The vertical line marks when the IAF model of the Giant Fiber SIZ emits a spike. At that time, the solid angle of a near-miss is nearly indistinguishable from a direct-hit’s, and separate only later. **(c)** Center elevation *ψ*(*t*) for the same trajectories. Here the traces have already fanned apart by the time the GF spikes for the direct-hit: the direct-hit stays on the line of sight (*ψ* 0) while the near-misses have drifted upward, in proportion to their pass-by distances. Thus elevation, not subtended solid angle, is what distinguishes the trajectory classes when the decision to initiate escape must be made.

As a result, the IAF neuron that integrates *z*(*t*) crosses the firing threshold only for the direct-hit. As shown in Figure 6b, at the moment the GF emits a spike for the direct-hit, the subtended solid angle for near-misses is almost identical to that of the direct-hit. Thus based on the solid angle alone one cannot distinguish near-misses before the GF spikes for the direct-hit, *i.e.,* it may be too late to make a decision based purely on the solid angle. As shown in Figure 6c, by the time the GF spikes for the direct-hit, the center elevation for near-misses has already grown in proportion to their pass-by distance. Thus center elevation distinguishes direct-hits from near-misses much earlier than the solid angle.

Therefore, we see that the subtended solid angle grows for direct-hits and near-misses alike during the early part of the approach, and falls for a near-miss only late, once the looming object has already passed the fly. A near-miss therefore looks much like a direct-hit in terms of solid angle during exactly the window in which the escape decision must be made. Our model, by contrast, responds not only to the increasing solid angle but also to the rising elevation of a near-miss’s center, which sets the two trajectory classes apart earlier, while there is still time to withhold the escape.

This places the Giant Fiber escape in its ethological setting, where our model enables timely and selective escape across the slow, close-range approaches that a fly can realistically evade, for the firing threshold chosen above. Beyond *v >* 0.5 m/s, the approaching object subtends a solid angle too small to be resolved by the few LPLC2 neurons that are active before the escape deadline, hence a fast aerial pursuer such as a dragonfly is effectively invisible until it is too late to escape. This is consistent with the near-total capture success that such predators achieve against flying prey [36].

In summary, we demonstrated that the circuit we proposed in Figure 4 separates direct-hits from near-misses by responding to the rising elevation of a near-miss’s center, well before the subtended solid angle could tell them apart.

## 4 Discussion

Connectomic datasets provide increasingly comprehensive reconstructions of neuronal morphology and synaptic connectivity, offering unprecedented opportunities to explore the structural organization of neural circuits. Abstracting these datasets into functional models remains a central challenge in neuroscience. Our connectome-grounded functional modeling results for the LPLC2 GF escape pathway show that while analyzing connectomic datasets reveals intriguing details of circuit architecture, exploring its functional logic requires two additional pieces of the puzzle: (i) modeling looming object trajectories in the visual space, and (ii) modeling the processing in each circuit and how they jointly give rise to collision detection.

Our first step was to formulate a looming object’s direct-hit or near-miss trajectories in physical 3D space, and their projected image on an ideal hemispherical eye, as illustrated in Figure 1a and in detail in Figure 10. Unlike a general visual scene, looming objects admit a compact and controlled description based on the retinal solid angle projection and its center elevation. This let us tie each model detection circuit output back to a well-defined looming object in the visual field and probe the circuit systematically across the range of approaches relevant to escape. Without such a handle on the visual field, the functional logic of the circuit, *i.e.*, the mapping from input to output that we seek to uncover, remains difficult to pin down.

Discriminating an object on a collision course from one that will pass by is a classic processing of insect looming detectors. In the locust, the descending contralateral movement detector is tightly tuned to collision trajectories, responding maximally to a direct approach and progressively less as the trajectory deviates toward a near miss, in both azimuth and elevation [14, 37]. Such selectivity is central to the collision-detection framework these circuits exemplify [38]. In *Drosophila*, by contrast, this discrimination has not been directly probed: the GF response is invariant to the azimuth of a centered looming stimulus [39], but its dependence on the elevation of the expansion center, and its response to non-colliding trajectories, remains untested. The looming object trajectory model allowed us to theoretically and computationally explore this question.

To examine the connectome-grounded functional logic of collision detection, our second step was to formulate how the collision-detection circuit, as depicted in Figure 4, represents and processes the looming object trajectory. In our model, each stage acts directly on the geometric quantities that describe the approaching looming object: the LPLC2 population represents the object’s solid angle as a retinotopic disk, and the GF reads that representation through the difference between its two dendritic branches, an operation tuned to the elevation of the expansion center, the very feature that separates a hit from a miss. Because both the representation and the processing are anchored in the visual field, the circuit’s functional logic becomes legible: a specific geometric property of the approaching object can be followed from the input through to the escape decision.

Our model provides a concrete, connectome-grounded hypothesis for how the fly performs this classic collision-detection, in which the GF’s escape input signal is processed by using the biphasic differential profile **C**_1_ **C**_2_ of its two dendritic branches. The input signal is inaccessible unless the two dendritic branches are resolved individually *and* each contributing axon is placed in columnar coordinates. This is precisely the combination of sub-neuronal and population analysis that NGB provides. As **C**_1_ **C**_2_ is azimuthally symmetric and tuned in elevation, and moreover consistent with the observed azimuthal invariance of the GF response [39] it suggests a specific, testable prediction along the elevation axis.

As in our connectome-grounded functional model, prior models of looming detection in *Drosophila* also incorporated a population of LPLC2-like units with dendritic subunits exhibiting radial motion opponency [17, 40]. In this work we characterized the geometric structure of the LPLC2 population code as an activation annulus, and additionally demonstrated that both the subtended solid angle and the center elevation can be estimated accurately from it. Furthermore, in these previous models the GF dendrite was modeled as a single unit, whereas in our model the two GF dendritic branches are modeled as separate units whose outputs are subsequently combined to drive the GF axon. It is this functional separation of the two dendritic branches that provides access to the object’s center elevation, which differentiates direct-hits from near-misses much earlier along the trajectory than the subtended solid angle alone. The solid angle stays nearly identical for direct-hits and near-misses until the object is too close to the fly, so a readout of the solid angle alone will force a decision between either escaping too early or waiting too long to escape, without leaving a middle ground of tradeoff between the two.

We point out that our model processes separately the cholinergic LPLC2 inputs into the two GF dendritic branches. It remains an open question whether the subtraction can be achieved between two dendritic branches with the same pre- and post-synaptic neuron pairs. While electrophysiological experiments showed that the GF is excited when all LPLC2s are optogenetically activated [27], it is unclear whether every individual LPLC2 neuron contributes equally as excitation, as such a whole-field activation pattern would never occur naturally. Note that, our model does not contradict the optogenetically activated electrophysiology results. A possible mechanism is a differential expression of acetylcholine receptors on each branch. Recently discovered inhibitory muscarinic acetylcholine receptors in the KCs [41] is a possible candidate for one of the branches while the other branch can express more typical, excitatory acetylcholine receptors. Regardless, as we have observed this biphasic profile in two different connectome datasets, our work suggests that sub-neuronal structures in a neuron may act more fundamentally as separate computational units. This calls for experiments with access to individual branches for a deeper understanding of the functional logic of fly brain circuits.

Arriving at this functional logic, however, was possible only because we could first recover the underlying pathway structure from the connectome. The NGB analysis primitives supported this recovery by providing easy, interactive access to both the sub-neuronal and the population-level structure of the pathway, and are applicable well beyond this looming detection circuit.

Connectomic reconstructions are growing rapidly in both size and complexity, extending from single neuropils to whole brains and increasingly spanning multiple datasets and modalities. Comprehending the structure of connectomic morphology at this scale, and turning it into functional understanding, outpaces what manual inspection or bespoke, hand-written analysis code can sustain. This is where the power of AI becomes essential. The contribution of NGB introduced in this work is to bring an AI Agent into the analysis workflow itself, where its role is to make the analysis *interactive* and to let it operate *at scale*. These are the two levers through which AI changes how connectomes can be studied.

A central benefit of introducing an AI Agent into connectome analysis is the interactivity it affords. Because the Human Explorer engages the workspace through natural-language dialogue, formulating questions and refining them through successive follow-ups, an investigation can proceed as an open-ended inquiry rather than a predetermined analysis: one need not decide in advance which analyses to perform or how to compose them, but may pursue, extend, or redirect a line of questioning as understanding develops. Where a question cannot be expressed in language alone, the Human Explorer can provide visual inputs on the canvas by direct manipulation of the 3D morphology, for instance selecting where to partition a dendrite or how to orient a projection manifold. This interaction closes the gap between visually inspecting a circuit’s morphology and analyzing it in natural language, allowing the agent to access the morphology in its natural 3D space. The significance of this interactivity is that it keeps the scientist’s own reasoning at the center of the analysis: hypotheses can be posed, tested, and reframed continuously, and the effort of exploration is spent on scientific judgment rather than on translating each question into code. In doing so, the workspace lowers the barrier to sub-neuronal circuit analysis and makes exploratory, hypothesis-driven investigation of the connectome a natural working modality.

A second benefit of introducing an AI Agent is the scale of exploration it makes possible. A circuit is rarely understood in isolation; exploring it requires bringing in more cell types, and comparing it across datasets. Each such extension adds further factors and structural aspects that must be held together within the same line of inquiry. Carried out by hand, this effort compounds with every element added, which is ultimately what limits how far an investigation can be taken. The value of scaling is therefore not merely performing the same analysis faster, but decoupling the reach of an inquiry from the effort it demands: the scope can grow, in the length of a pathway, the number of cell types, and the number of datasets, while the work of orchestrating it is automated rather than borne by the Human Explorer. Retaining the state of an investigation and composing the analysis primitives on demand are the means by which this is accomplished: a single line of inquiry can extend across many turns and reach across neuropils and datasets, so that a circuit can be examined in its brain-wide context across different datasets through a common interface (see also Section 2.4). It is worth emphasizing that this scaling is not an immediate consequence of the language model itself, but a result of the deliberate design of the underlying analysis API and the curated Model Context Protocol interface through which the primitives are exposed (see also Appendix A.1): the agent supplies the natural-language front end, whereas the engineered toolset, with its typed primitives, composer functions, and cached derived quantities, is what actually delivers the scale.

The significance of working at scale enables the study of the connectome-grounded functional logic of looming circuits. Recovering the functional logic of a circuit, as for the looming pathway above, requires holding the circuit in view as a whole, its sub-neuronal structure, the arrangement of its population, and the way these together transform input into output, since this logic resides in the organization of the whole rather than in any single neuron. Experimental approaches that record from only one neuron or a handful at a time cannot assemble a picture of this kind, and the functional logic of a large circuit accordingly remains difficult to probe experimentally. Analyzing the connectome at scale lifts this constraint: it allows the functional logic to be explored across increasingly comprehensive circuits, so that scaling the analysis is, in effect, scaling the study of functional logic itself.

More broadly, the NGB workspace is positioned to serve as a unified interface to the wider ecosystem of connectome-analysis tools, exposing each through the same agentic, natural-language front end that are separately implemented in the polysynaptic path-finding of the Connectome Interpreter Toolkit [42], the code-free single-dataset exploration of the FlyWire Codex, the custom per-dataset pipelines behind brain-scale pathway analyses [43], and the expert-curated neuroanatomical ontology that VFB-MCP exposes from Virtual Fly Brain [44]. Composed alongside existing NGB primitives of sub-neuronal segment structure, population topography, and retinotopy, such tools can be invoked and combined interactively across spatial scales and datasets from within a single workspace.

The collision-detection circuit we modeled in this paper illustrates the workflow for exploring the connectome at scale to uncover its connectome-grounded functional logic. It is not, however, a complete account of the circuit, and several directions remain open. On the input side, the trajectory model of the looming object moving in the visual field could be enriched: rather than holding the azimuth fixed, near misses could be given time-dependent azimuth values. Head turns driven by the Central Complex could also be incorporated to actively control the azimuth. On the circuit side, our model is purely feedforward, yet feedback components such as the LPi cell types in the LOP are present at almost every stage of visual processing. The LPLC2s also receive synaptic inputs from additional cell types in the Lobula (LOB, and we hypothesize that they could provide additional spatiotemporal information during head turns and saccades [45]. The LC4 pathway [16, 46] provides a second major input to the GF, and is also primarily driven by Tm inputs in the LOB. Incorporating these structures, together with a richer model of trajectories, could extend the account beyond elevation-based discrimination and reveal how feedback and parallel input pathways jointly shape the GF’s escape decision. Pursuing these directions within the same interactive, scalable workflow is precisely what NGB is designed to support.

Taken together, our work provides a complete workflow that carries a neural circuit from morphology, through a pathway abstraction, to an executable connectome-grounded functional model that generates concrete, experimentally testable predictions. NGB is designed to make this workflow repeatable: the same primitives, agentic workspace, and pathway abstractions that we applied to the LPLC2 GF escape circuit apply to other neural circuits throughout the fly brain, and we expect NGB to accelerate the translation of connectomic structure into circuit function, well beyond the visual escape pathway studied here.

## A NeuroGraphBench: Architecture, Analysis Primitives, and the Interactive Workspace

### A.1 NGB Architecture

The NeuroGraphBench backend architecture consists of a three-layer server and database stack schematically depicted at the bottom of Figure 3. *NeuroArch* [6] is a graph database that stores, under a unified schema, neuropil meshes, neuron morphology (both surface meshes and tree-structured skeletons), synapse tables with pre/post neuron and celltype annotations, and neurotransmitter predictions across four Drosophila connectome datasets: OpticLobe [3], Hemibrain [1], FlyWire [2], and L1EM [4]. Each connectome dataset is loaded as a separate *Neo4j* instance.

Above the NeuroArch database, is an API server called *FlyBrainAtlas* (FBA). Built with Python and FastAPI, FBA forwards raw graph queries to the Neo4j instances and serves their results, along with precomputed derived quantities held in an on-disk cache. The cache is organized along NGB primitives (see Section 2.1). For *volumetric segmentation*, per-neuropil synapse tables tag every synapse with its containing neuropil boundary to support neuropil-scoped connectivity queries. For *skeletal segmentation*, per-neuron synaptic tables index each synapse to its nearest skeleton node to support local neuron connectivity queries. For *columnar mapping*, curated hex-column indices pin every columnar neuron to a discrete hexagonal coordinate. As an example, an index derived from the topography of L1, L2 and Mi1, Tm1, respectively, anchors the lamina and medulla columnar grids, and per-family lookups propagate the coordinates along the ON (Mi1*→*T4) and OFF (Tm1*→*T5) motion pathways. For *topographic mapping* no persistent cache is needed, because the primitive is anchored by an interactively placed projection manifold (a two-dimensional surface), and morphology data is streamed from NeuroArch on demand. A separately cached *celltype connectivity graph*, built by aggregating per-neuropil synaptic tables into (pre-celltype, post-celltype, neuropil, layer, count) tuples, and a companion *tract graph* of directed celltype-to-celltype summaries with polarity classification, together support the pathway-search described below.

Above FBA, the *NeuroGraphBench Client* wraps the FBA APIs into typed Python functions and exposes them as tools to an AI Agent through a Model Context Protocol (MCP) server. We used *Claude Opus 4.8* for the demonstrations in this manuscript, noting that any other LLM model that supports MCPs can be used. Crucially to ensure correctness, the AI agent is not allowed to write any new code and is entirely restricted to only invoke FBA API calls.

Frontend rendering runs on four complementary canvases: (i) a 3D canvas built on the *neu3D* library (https://github.com/FruitFlyBrain/neu3d) that interactively visualizes neuropil meshes, neuron skeletons and meshes, synapse point clouds, and controls for topographic-manifold placement and skeletal-anchor selection, (ii) a Connectivity canvas built on *cytoscape.js* [47] that draws celltype-level pathway diagrams with nesting by neuropil and layer, (iii) a Retinotopy canvas that draws SVG on the hex-column lattice, and (iv) a Topography canvas that draws SVG scatter plots of projected neuron locations on user-placed manifolds. Each canvas maintains its own persistent state on the server, and both the Human Explorer and the AI Agent act on it by invoking the same backend commands; the resulting updates stream to the frontend over a per-canvas WebSocket channel.

The Human Explorer and the AI Agent drive these canvases through two parallel channels. Through the chat panel, the AI Agent parses a natural-language question into a sequence of typed MCP calls, each dispatched through the FBA server and served from the cache or forwarded to Neo4j instances; the result is committed at once to the chat panel as a tool-grounded answer, to the appropriate canvas as a scene mutation, and to a persistent *workspace state* that tracks canvas elements, findings, and open questions across turns. This persistence turns a sequence of one-shot queries into a cumulative investigation. For example, a neuronal segment (such as an axonal or dendritic branch) created several turns earlier (via volumetric or skeletal segmentation) remains addressable by its short identifier (e.g., segment 38472), so that the AI Agent, asked about “the LPLC2 segment I created,” consults the workspace state rather than reconstructing it from memory.

Through the three-dimensional canvas, the Human Explorer interacts with the morphology directly: configuring the geodesic neighborhood for skeletal segmentation and the projection manifold for topographic mapping, and copying the name of any element on the canvas to then chat with the AI Agent about it. These selections pass to the server as typed WebSocket messages and enter the workspace state, so that the agent can reason about them in later turns.

NGB also supports the export of an executable *provenance record* for a session containing the natural-language queries, every typed tool call with its parameters, and the geometric anchors supplied by the Human Explorer. This enables direct replication in an independent Python environment outside the NGB workspace, using only the FBA API. Figure 7 shows an example of such a provenance record for a short session in the NGB workspace.

**Figure 7:**
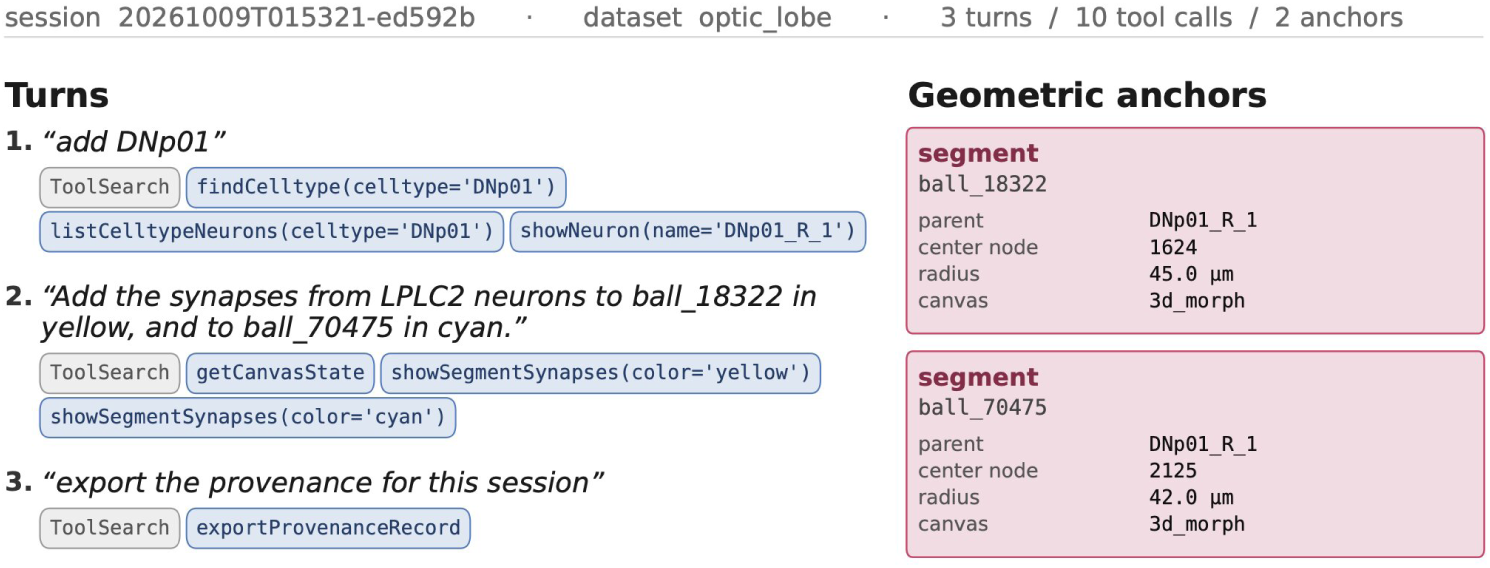
Example of a provenance record exported from the NGB workspace. The record header contains the session id (20261009T015321-ed592b), the connectome dataset it was run against (optic lobe), and the size of the record: three turns, ten tool calls and two geometric anchors. The record also lists, for each turn of the session, the Human Explorer’s natural-language query followed by the FlyBrainAtlas API calls that the AI Agent invoked to answer it, along with their arguments; the grey chip marks the agent’s lookup of the available tools. In the first turn the agent resolves the cell type DNp01 and renders the neuron, in the second it adds the LPLC2 synapses onto two previously created segments, and in the third it exports the record itself. The record also stores geometric anchors that the Human Explorer supplied by direct manipulation on the 3D canvas: here two geodesic balls on the skeleton of DNp01 R 1, each recorded with its parent neuron, center node, radius and the canvas on which it was placed.

### A.2 NGB Analysis Primitives

Neurons are embedded in the 3D volume of the brain; from this perspective, brain morphology is characterized by the extrinsic geometry of dendritic and axonal arbors occupying anatomically defined regions such as neuropils and layers. Complementing this extrinsic view, each neuron also possesses an intrinsic organization, independent of its 3D embedding, defined by the tree-like topology of its skeletal segments. Above the level of a single neuron, populations of neurons of the same type frequently exhibit emergent spatial organization, such as parallel bundles of neurites, columnar tilings, and columnar lattices, that are not captured by any single-neuron description. The four NGB analysis primitives (see Figure 8) are designed to provide programmatic access to this sub-neuronal structure and spatial organization of neural circuits in the fly brain.

**Figure 8:**
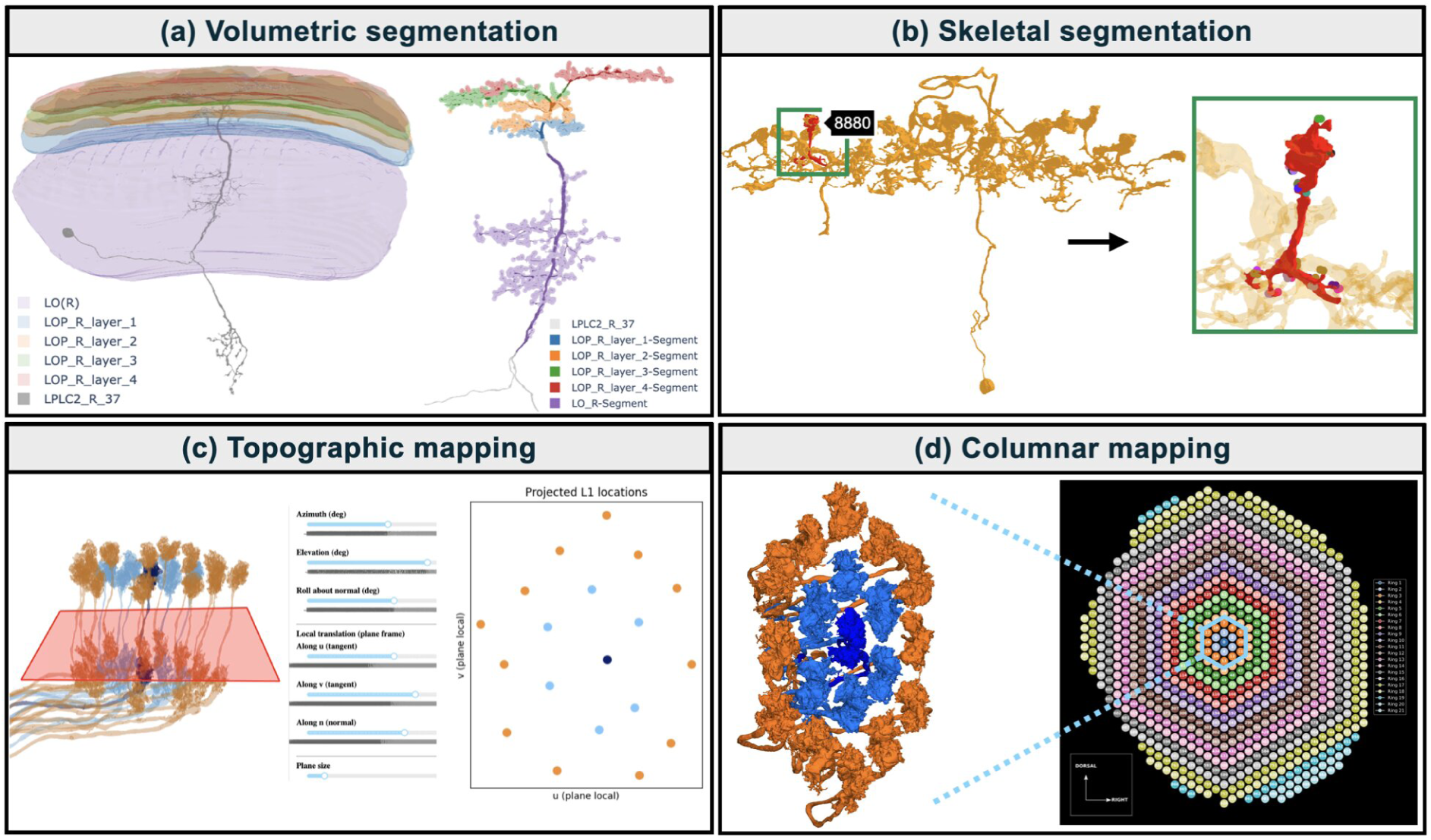
The four NGB analysis primitives. **(a)** *Volumetric segmentation:* (left) an LPLC2 neuron (gray) with dendritic arbors innervating four LOP layers (LOP-1…4, colored) and the lobula (LO); (right) the neuron is volumetrically segmented using the neuropil/layer boundary, with neuron segments and corresponding synapse locations colored in each neuropil/layer. **(b)** *Skeletal segmentation:* hovering near an LPi34 neuron (yellow) displays the database ID of the closest neurite fragment along its skeleton (here, 8880); a local geodesic neighborhood along the skeleton (red), centered at the chosen neurite is retrieved together with the synapses arborizing within it. **(c)** *Topographic mapping:* 19 neighboring L1 neurons and a user-placed planar projection manifold (red); the interface provides controls to translate, rotate, and resize the manifold relative to the neurons; intersection locations of each neuron with the manifold are shown at right, arranged approximately in a hexagonal grid. **(d)** *Columnar mapping:* 19 neighboring L1 neurons arranged in three concentric rings, retrieved by specifying the set of retinotopically organized columns they innervate in the lamina; locations of columnar neurons in the optic lobe were mapped to points on a hexagonal grid comprising 890 columns arranged in 21 concentric rings.

**Volumetric segmentation** partitions a neuron by the regions it innervates and retrieves the arbors and synapses that fall within each partition (Figure 8a). A *region* is a volumetric subset of the brain specified by a closed surface mesh; a *subregion* is a region whose mesh is fully contained within a parent region, such as a layer within a neuropil.

**Skeletal segmentation** extracts a local neuronal segment of a single neurite as a *geodesic neighborhood* on the neuron’s skeleton: a user-selected anchor point serves as the center of the neighborhood, which comprises the neurites within a chosen shortest-path distance of the anchor along the skeleton, together with the synapses located on them (Figure 8b). The anchor is selected visually on the 3D canvas, since a specific neurite on a large skeleton is hard to address by a database identifier alone.

**Topographic mapping** assigns continuous 2D coordinates to a population of neurons by intersecting each neuron’s morphology with a user-placed *projection manifold*, a 2D surface embedded in three-dimensional space (Figure 8c). The interactive placement fixes the coordinate system for the analysis and requires the Human Explorer to view the neuronal tract in 3D before choosing an appropriate surface.

**Columnar mapping** projects columnar-neuron locations onto a discrete hexagonal coordinate system of the ommatidia/cartridge/columns, and propagates the coordinates along one-to-one feedforward pathways in the ON and OFF motion channels (Figure 8d). For a non-columnar neuron, its columnar coverage is the set of labels of its columnar synaptic partners, and represents the region of the visual space it operates in. The estimate of the region where the LPLC2 neurons are located, for example, is inferred from the locations of their T4 and T5 inputs (see also Section 3.1).

We applied the NGB analysis primitives to the LPLC2 GF pathway in the OpticLobe and Hemibrain datasets in Section 2.1 and Section 2.2. In particular, skeletal segmentation and topographic mapping were performed on the Hemibrain dataset that has incomplete reconstructions of the LPLC2 dendrites and on the Optic Lobe dataset that prevents the use of columnar mapping. To further corroborate our findings from skeletal segmentation of the lateral GF dendrite and to provide a direct comparison between columnar mapping and topographic mapping, we repeated this analysis in the MaleCNS dataset as shown in Figure 9.

**Figure 9:**
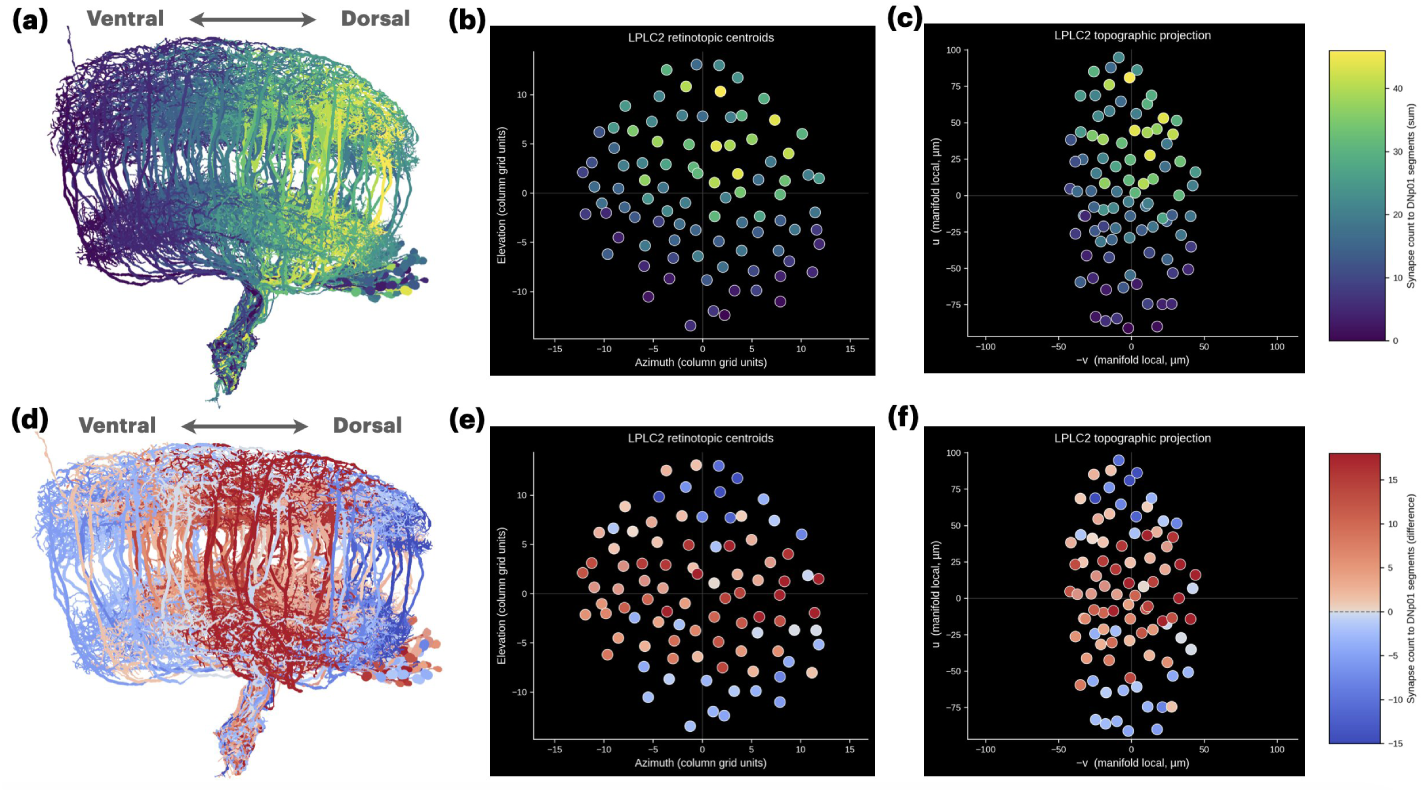
Spatial profiles of LPLC2→GF synapses in the MaleCNS dataset. The sum **C**_1_ + **C**_2_ and difference **C_1_ C_2_** of the synaptic counts from each LPLC2 to the dorsal and ventral branches of the lateral GF dendrite, respectively visualized by coloring in **(a-c)** and **(d-f)**: **(a, d)** the LPLC2 population morphology, **(b, e)** LPLC2 centers estimated from columnar T4 and T5 inputs, and **(c, f)** topographically estimated LPLC2 axon locations (in the Lobula).

We observe the same spatial profile for synaptic counts from the LPLC2 population to the dorsal and ventral branches of the GF dendrite. The columnar and topographic mapping are visually consistent. Here we note that the topographic map is compressed along the anterioposterior axis due to the corresponding flattened shape of the Lobula (where the projection manifold was placed to intersect the LPLC2 axons, as in Figure 2j).

## B Modeling the Looming Object Trajectory and the Looming Detection Circuit

This section gives the full theoretical grounding of the executable circuit model shown in Figure 4 in Section 3.1.

### Modeling the Looming Object Trajectory

Before describing the looming detection circuit, we model the trajectory in the visual space of a looming object approaching the fruit fly retina. We model the retina as a hexagonal lattice of 890 ommatidia tiling the surface of a unit sphere, arranged in 21 concentric rings with a uniform interommatidial angle of 4.5*^◦^* (see also Figure 8d). The luminance of the 3D object is sampled along the optical axis of every ommatidium, and the resulting image is the input to the retina.

#### Projection Geometry

We define a spherical coordinate system with the fly eye at its origin. Then a spherical object of radius *r* at radial distance *d*(*t*), azimuth *ϕ*(*t*), and elevation *ψ*(*t*) (measured from the line of sight) subtends an angular diameter and a solid angle

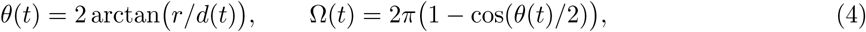

each a function of *d*(*t*) alone for a fixed object size. We restrict the looming objects to trajectories that drift in elevation, holding the azimuth at *ϕ* = 0 and parameterizing each trajectory by its radial distance *d*(*t*) and elevation *ψ*(*t*). The choice of *ϕ* = 0 is a deliberate restriction to the behaviorally relevant dorsoventral near misses along the elevation axis. A purely azimuthal drift is a distinct case not considered here, and a combined up-and-sideways trajectory would separate along a shallower effective-elevation path.

#### Object trajectories

Under the assumptions above, every trajectory is specified by two quantities: the approach velocity *v* and the pass-by distance *b*, the object’s closest radial distance to the fly’s eye. A single object radius *r* = 0.01 m (a 2 cm diameter, matching the frontal cross-section of small aerial predators such as damselflies and robber flies) is used throughout. As shown in Figure 10, the object travels in a straight line that begins on the line of sight at a distance of *d*(0) = 25 cm from the fly and passes the fly at closest radial distance *b*, approaching at velocity *v*; if the time at the closest approach is *t*_ca_, its radial distance and elevation are

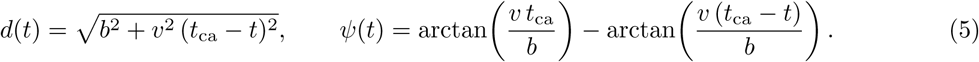

**Figure 10:**
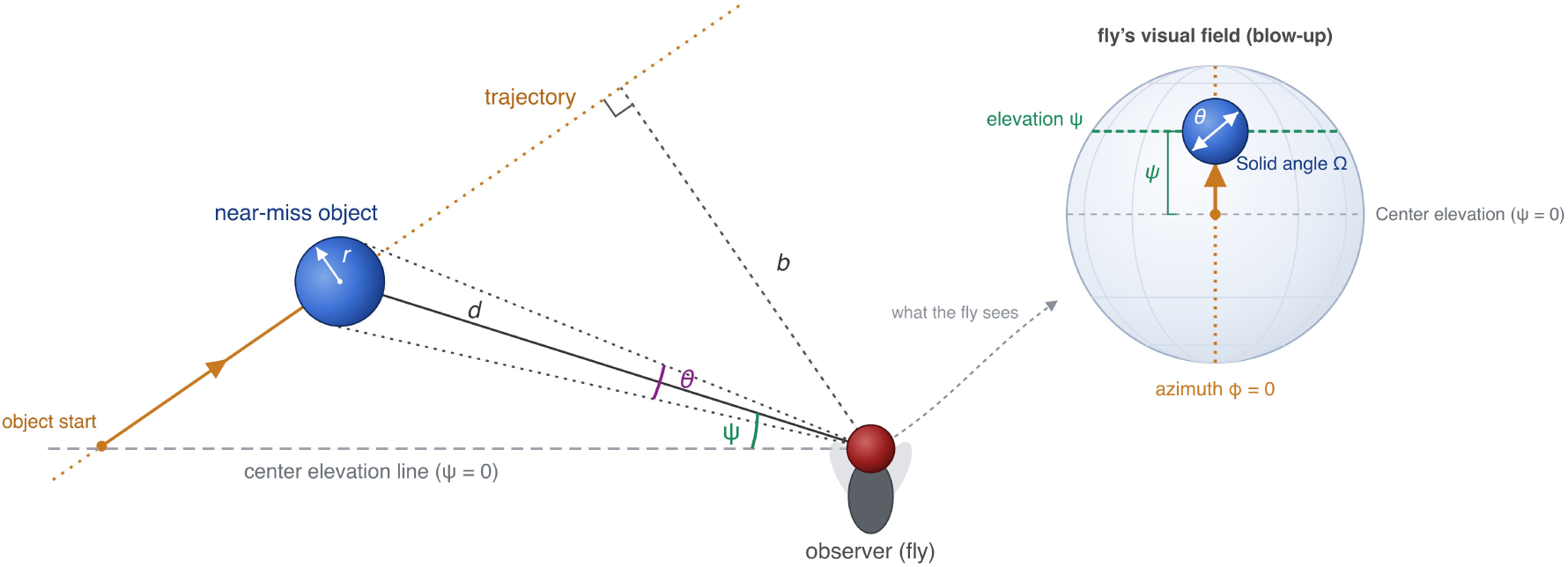
Geometry of the trajectory of a spherical visual object for a near-miss scenario. An object of radius *r* (blue sphere) approaches the fly (drawn with its spherical eye, upright and facing up) along a straight-line trajectory (yellow dotted) that begins on the center elevation line (*ψ* = 0, the line of sight) and drifts upward. At the instant shown it lies at radial distance *d* and elevation *ψ* above the line of sight, subtending an angular diameter *θ* (dotted rays to the object’s edges) and solid angle Ω = 2*π* 1 cos(*θ/*2) . Its closest approach to the fly amounts to the distance *b*, with the right angle marked. The blow-up on the right shows, at the same instant, the projection of the visual object onto the fly’s spherical eye: the object’s image lies on the azimuth *ϕ* = 0 (orange), and moves along it as its elevation *ψ* increases (green line through the image center), so that only the elevation varies along the trajectory; the image subtends the angular diameter *θ* (white).

The angular diameter *θ*(*t*) = 2 arctan *r/d*(*t*) rises to a maximum 2 arctan(*r/b*) at closest approach and then falls as the object recedes; the elevation *ψ*(*t*) starts at 0 and sweeps upward as the object passes, and for *t > t*_ca_ the object goes behind the fly and leaves the frontal field. Eq. (5) describes both types of trajectories, which differ only in the range of *b*: a *direct-hit* (DH) has a small pass-by distance, *b* 1 mm, so the object strikes the fly and its expansion stays essentially centered; a *near-miss* (NM) has *b >* 1 mm, so the object center drifts out of the field of view. For *b* = 0 the trajectory reduces to *ψ*(*t*) *≡* 0 with *d*(*t*) = *v* (*t*_ca_ *− t*).

### Modeling the Looming Detection Circuit

With time denoted by *t*, the axonal outputs of T4 neuron subtypes (T4a, T4b, T4c, T4d) and T5 neuron subtypes (T5a, T5b, T5c, T5d) in column *j* (for *j* = 1*, . . . , M* ) are respectively denoted as 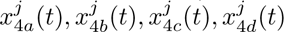 and 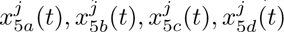. Each of these T4/T5 subtypes is modeled as a three-arm elementary motion detector [48, 49].

The motion signals 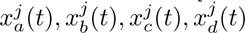, indexed by subtypes (*a, b, c, d*), denote the sum of outputs from the respective T4 and T5 subtypes, and are estimates of local motion along the left, right, up and down directions respectively, for the modeled right eye as defined in Section 2.1. The motion signal of each subtype is the sum of the outputs of the corresponding T4 and T5 cells in that column,

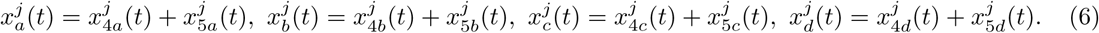

Together, these motion signals define a retinotopic *motion field* that serves as input to the looming detection circuit.

There are *N* LPLC2 neurons, with the center of neuron *i* denoted by *P ^i^*, for *i* = 1*, . . . , N* . The four dendritic subunits of each neuron *i* are modeled as spatial filters with entries 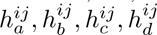 in column *j* that respectively process (*a, b, c, d*) motion signals from T4s and T5s neurons. Each filter features an elliptical kernel oriented along its preferred direction, and the four filters share *P ^i^* as a common center, producing the radial arrangement of Figure 11a (for more details, see the connectomic morphology of Figure 2a-c). The LPi interneurons are modeled as providing opponent inhibition, so that each subunit’s drive is the difference between motion in its preferred direction and motion in the opposite direction (Figure 4a). Note however that, motion in the two opposite directions cannot coexist in the same column. Therefore, in each column the subunit receives either a excitatory drive from motion in the preferred direction or a inhibitory drive from motion in the opposite direction. Each dendritic subunit’s output is a weighted sum of the motion fields over the columns, and the LPLC2 axonal output *y^i^*(*t*) is obtained by summing the four subunit outputs, followed by half-wave rectification as,

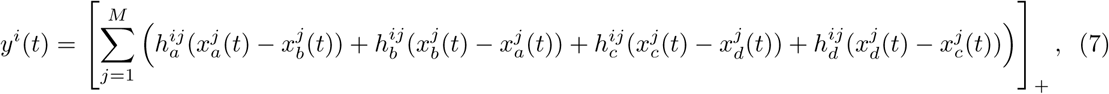

where[ ]_+_ is the half-wave rectification.

**Figure 11:**
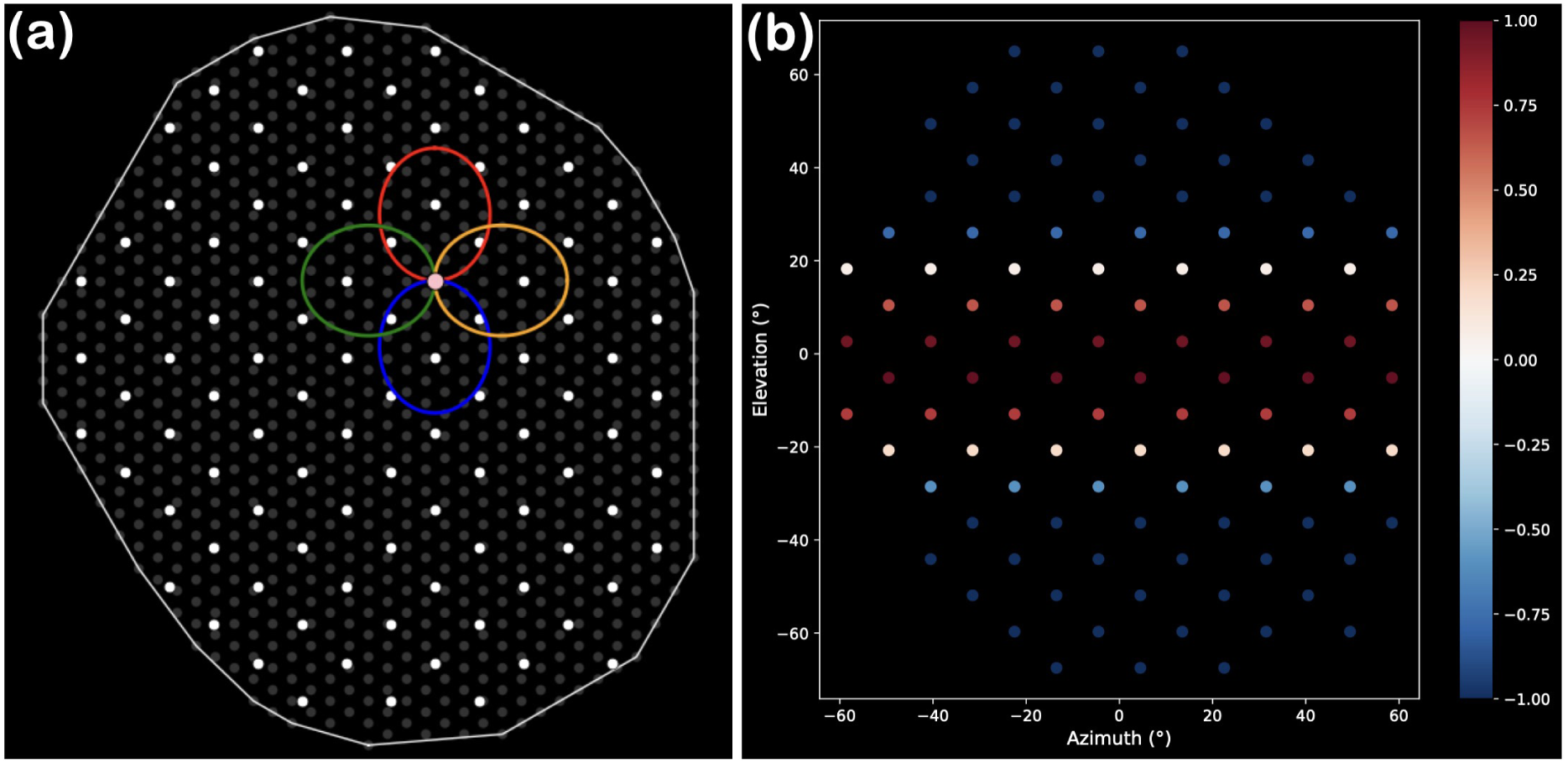
Models of the LPLC2 dendritic arbors and the Giant Fiber synaptic weight vector C_1_ C_2_. **(a)** Centers (white dots) of the *N* LPLC2 neurons. One LPLC2 center is highlighted in pink, with its associated dendritic arbors modeled as spatial filters for motion in the left, right, up, and down direction depicted in green, orange, red, and blue, respectively. **(b)** the Giant Fiber synaptic weight vector **C**_1_ **C**_2_ modeled as a quadratic function of elevation, positive within a band centered on the population mean elevation and negative outside it. The LPLC2 centers are colored by their corresponding weights (red positive, blue negative).

The Giant Fiber reads the LPLC2 population **y**(*t*) through the two sub-branches of its lateral dendrite (see also Section 2.1), each modeled as a linear weighted sum:

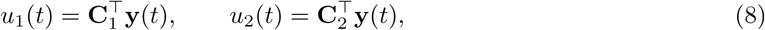

where **C**_1_ (dorsal sub-branch) and **C**_2_ (ventral sub-branch) are nonnegative weight vectors, and **y**(*t*) = *y*^1^(*t*)*, y*^2^(*t*)*, . . . , y^N^* (*t*) *^⊤^* is the LPLC2 population vector.

Finally, the input to the Giant Fiber SIZ in Figure 4 amounts to

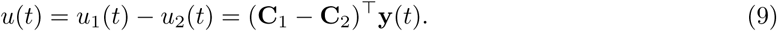

We modeled the difference of the weight vectors **C**_1_ **C**_2_ as a quadratic function, whose value for the LPLC2 neuron *i* with center *P ^i^* and elevation *ψ^i^* (measured from the center elevation line) amounts to

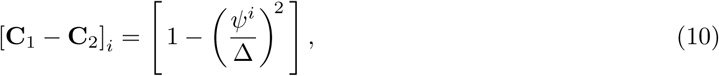

where Δ = 21*^◦^* so the weights are positive in the elevation band with width 2Δ = 42*^◦^* and negative otherwise, in line with connectomic findings (see Figure 2i,l and Figure 9d-f). Note that the values of the weight entries do not depend on the azimuth.

The Giant Fiber SIZ converts the differential Giant Fiber input *u*(*t*) into an escape decision in two stages. First, a divisive normalization processor (DNP) normalizes *u* by the square of its own temporal derivative (see also Figure 4)

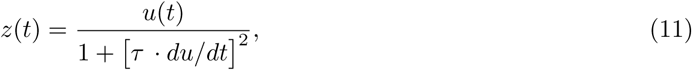

where *τ* = 20 ms is a time constant. Second, the DNP output *z*(*t*) is passed to an integrate-and-fire (IAF) neuron, that generates an escape spike the first time its membrane potential reaches threshold *δ*.

## C Computational Evaluation of the Looming Detection Circuit

This section provides additional computational insights gained while exploring the functional logic of the looming detection circuit detailed in Section 3.2. In particular, we show that the solid angle and elevation, that we used to distinguish direct-hits and near-misses in our model, can be estimated accurately from the modeled LPLC2 population activity.

As shown in Figure 12a, the centers of the LPLC2 neurons tile the visual space along a hexagonal grid with angular spacing *s* = 12*^◦^* between neighboring LPLC2 centers. See also Figure 11(a), where the white dots depict the LPLC2 centers used in our simulation. A looming object of uniform luminance generates local motion only at the boundary of the retinal disk *D*(*t*) it subtends, as its interior and the surrounding background are both uniform for the explored direct-hit and near-miss trajectories. An LPLC2 receives motion inputs only if its dendritic receptive fields intersect the expanding boundary. The expanding boundary of the solid angle therefore drives the LPLC2s located in an annulus around it, where the width of the annulus is set by the radial extent of the LPLC2 receptive fields (27*^◦^* in our simulations, the length of the major axis of a dendritic subunit measured from the shared center).

**Figure 12:**
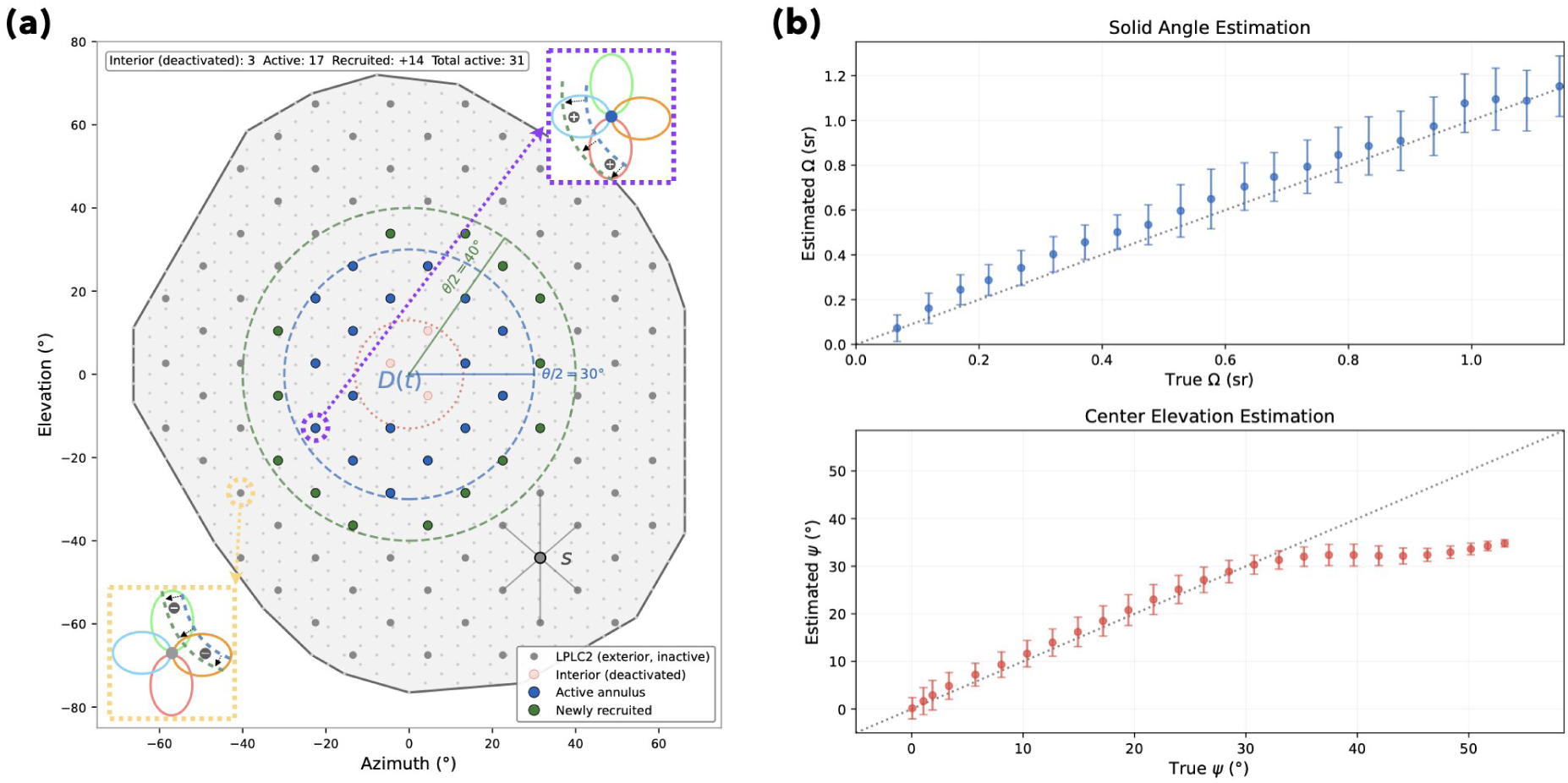
Estimation of the solid angle and elevation given the activated population of LPLC2s. **(a)** *The annulus of activated LPLC2 neurons around the expanding boundary of the solid angle subtended by the retinal disk* (*t*). The LPLC2 centers lie on a hexagonal lattice with angular spacing *s* (the mean angular separation between neighboring centers). The expanding boundary of the subtended solid angle, of angular radius *θ/*2, drives the LPLC2s whose dendritic receptive fields it intersects. Of these, the neurons whose centers lie inside the subtended solid angle are activated (blue), while those whose centers lie outside remain inactive (grey dots). The purple and yellow insets depict the dendritic receptive fields intersected by the expanding disk boundary respectively for an LPLC2 in the interior and outside the subtended solid angle. In the purple inset, the boundary moves away from the neuron’s center, matching the preferred direction of the LPLC2 receptive fields, and the intersected receptive fields receive excitatory inputs from T4s and T5s. In the yellow inset, the boundary moves towards the neuron’s center and the intersected receptive fields receive inhibitory inputs from LPis tuned to the opposite direction, so that the rectified sum is zero. As the object grows the boundary expands (here from *θ/*2 = 30*^◦^* to 40*^◦^*, dashed circles), and the activated LPLC2 annulus moves outward with it, recruiting the neurons it reaches (green dots) while those it has passed are deactivated (pink dots). **(b)** *Solid angle and center elevation estimates from LPLC2 population activity.* Estimates are evaluated across 500 looming trajectories, simulated at 10 approach speeds (*v* = 0.10–0.55 m/s) and 50 pass-by distances (*b* = 1–50 mm), restricted to the approach phase before the disk boundary reaches the edge of the LPLC2 population. (Top) Estimated versus true solid angle Ω; the estimate closely tracks the identity line (dotted), with variance increasing for larger Ω. (Bottom) Estimated versus true center elevation *ψ*; the estimate is accurate for *ψ* ≲ 35*^◦^* and introduces larger error for larger elevations due to the finite retinal coverage of the LPLC2 population ( 65*^◦^*). Error bars show 1*^◦^* standard deviation across trajectories.

We distinguish two cases for an LPLC2 neuron with its center in the annulus that contains the solid angle boundary: (i) If the center is in the interior of the solid angle, the expanding boundary matches the preferred direction of the receptive fields it intersects (purple inset in Figure 12a), resulting in positive outputs that are retained after rectification. Thus this LPLC2 neuron is activated. (ii) If the center is in the exterior of the solid angle, the expanding boundary moves against the preferred direction of the receptive fields it intersects (yellow inset in Figure 12a). This results in negative outputs from these receptive fields. Therefore, the LPLC2 remains silent.

The activated set of LPLC2s is thus an annulus that grows with the expanding boundary of the subtended solid angle (Figure 12a). Its outer radius is the angular radius *θ*(*t*)*/*2 of the retinal disk and its centroid is the disk’s center elevation *ψ*(*t*), so the annulus traces the geometry of the solid angle. For a near-miss, in addition, the upper boundary expands faster than the lower one, so the annulus is activated asymmetrically, more strongly at higher elevations.

We note that a filled disk is a special case of the activated LPLC2 annulus: while *θ/*2 remains below the radial extent of the receptive fields, every LPLC2 inside the disk still intersects the expanding boundary, and so the active set is a solid disk. As the disk expands further, the central neurons fall out of reach of the boundary and hence the active set becomes a true annulus (see Figure 5b).

We proceed to demonstrate that the full retinal disk can be recovered from the activated LPLC2 annulus, by applying two processing steps. First, the activation of each LPLC2 is normalized by the maximum activation across the population at each time step, which removes the activation asymmetry between the faster and slower expanding sides of the boundary for near-misses.

Second, the normalized activations are processed by a nonlinear temporal filter: a temporal high-pass filter that boosts activation onsets at the expanding disk boundary, followed by a compressive power-law nonlinearity (with exponent 0.3), and a low-pass filter with time constant *τ* = 500 ms that holds the activation of each LPLC2 after the boundary has passed. As a result, LPLC2s that were activated when the expanding boundary swept through their receptive field retain their activation as the boundary moves outward, filling in the interior of the disk. Together, these steps recover a full, symmetric solid angle from the annular and asymmetric raw activity pattern of the population of active LPLC2 neurons.

To validate that the processed LPLC2 activation annulus faithfully represents the retinal disk *D*(*t*), we estimated the subtended solid angle and center elevation as follows: Since the LPLC2 centers tile the visual field, the sum of the processed LPLC2 activity multiplied with the solid angle subtended by a hexagonal tile (equal to 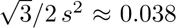 sr for *s* = 12*^◦^* expressed in radians) provides an estimate of Ω(*t*). The elevation *ψ*(*t*) of the retinal disk’s center is estimated as the activity-weighted mean of the elevations of the LPLC2 centers.

Figure 12b shows that the solid angle and center elevation estimates, evaluated across 500 looming trajectories spanning 10 approach speeds and 50 pass-by distances, are accurate except at high elevation primarily due to the fact that the model eye only covers 65*^◦^* of the visual space.

In summary, the size of the retinal disk , i.e., the subtended solid angle, and the elevation of the disk are faithfully represented by the processed LPLC2 activation annulus.

## Acknowledgement

The research reported here was supported, in part, by the National Science Foundation under grant #2400687.

